# Molecular basis of indispensable accuracy of mammalian miRNA biogenesis

**DOI:** 10.1101/2022.04.13.488181

**Authors:** David Zapletal, Eliska Taborska, Josef Pasulka, Radek Malik, Karel Kubicek, Martina Zanova, Christian Much, Marek Sebesta, Valeria Buccheri, Filip Horvat, Irena Jenickova, Michaela Prochazkova, Jan Prochazka, Matyas Pinkas, Jiri Novacek, Diego F. Joseph, Radislav Sedlacek, Dónal O’Carroll, Richard Stefl, Petr Svoboda

## Abstract

Mammalian Dicer is the gatekeeper into the essential gene-regulating miRNA pathway. What is committing mammalian Dicer to the miRNA pathway remains unknown. We report that Dicer’s highly conserved DExD/H helicase domain is the key structural element supporting accurate miRNA biogenesis. While ATPase activity of the domain is non-essential, its loss is lethal in mice. It is required during canonical miRNA biogenesis for efficient recognition, high-fidelity cleavage, and strand selection. Structure of Dicer-miRNA precursor complexes showed that the DExD/H domain acquired helicase-unrelated function defining Dicer conformations, which affect substrate loading and facilitate pre-selection of miRNA precursors. Dicer lacking the DExD/H domain favors conformations enabling reduced substrate selectivity and supporting RNA interference, a different small RNA pathway. Therefore, Dicer’s DExD/H domain ensures indispensable high-fidelity precursor processing of mammalian miRNAs, which constitutes a structural “mold” for adaptive miRNA evolution.

## Main Text

Dicer endoribonucleases generate small RNAs for RNA interference (RNAi) and microRNA (miRNA) pathways [1]. Dicer produces gene-regulating miRNAs by a single cleavage of genome-encoded small stem-loop precursors (pre-miRNA) [2], whereas processive cleavage of long dsRNA by Dicer gives rise to siRNAs that have gene-regulating or defensive roles against viruses or retrotransposons [3]. Vertebrate genomes carry a single highly conserved Dicer (*Dicer-1*) gene [4], which encodes ∼220 kDa multidomain protein and appears dedicated to the miRNA pathway. The N-terminal helicase domain and its activity are linked to evolution of specialized Dicer variants and divergence of small RNA pathways. This domain is composed of three globular subdomains: an N-terminal DExD/H domain (HEL1), which is separated by an insertion domain (HEL2i) from a helicase superfamily c-terminal domain (HEL2) (Fig. 1A). DCR-1 from *C. elegans*, Dicer-2 from *Drosophila*, and DCL1 *Arabidopsis* hydrolyze ATP [5-7], which enables threading dsRNA substrates through the helicase [8, 9]. The miRNA-producing Dicer-1 in *Drosophila* does not hydrolyze ATP and its helicase domain is degenerated [10]. Mice are unique among mammals because they employ two functionally distinct Dicers: the full-length Dicer for miRNA biogenesis and the oocyte-specific Dicer^O^ isoform, which lacks N-terminal DExD/H helicase subdomain (HEL1) and supports both, miRNA biogenesis and highly active RNAi[11]. At the same time, the highly conserved HEL1 in the miRNA-producing mammalian Dicer encompasses residues necessary for ATPase activity [4, 12] but it inhibits RNAi [13] and does not exhibit ATPase activity [14, 15]. Thus, the role and importance of Dicer’s helicase domain within endogenous mammalian small RNA pathways remains unknown.

**Figure 1.**
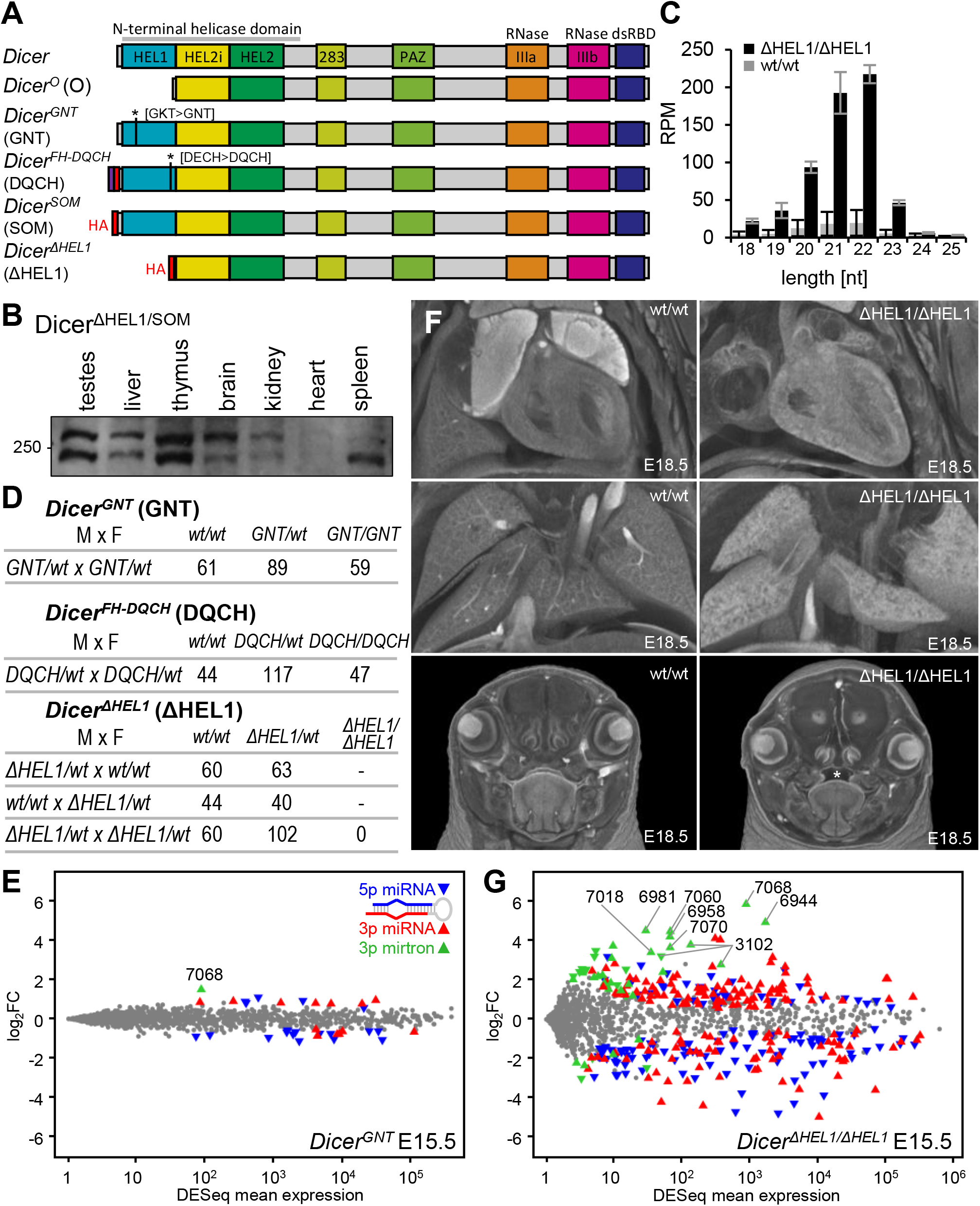
DExD/HEL1 domain of Dicer but not its enzymatic activity is essential for miRNA homeostasis and normal mouse development. (A) Schematic depiction of Dicer protein variants and mutants. On top are schemes of endogenous isoforms from mice: the full-length Dicer and Dicer^O^ lacking HEL1. Next are point mutations in HEL1 abolishing its function as ATPase and HA-tagged mutants with remodeled region encoding the N-terminus. We designated the engineered allele *Dicer*^Δ*HEL1*^ to distinguish it from the endogenous *Dicer*^*O*^ isoform, which is transcribed from a different promoter, has a different 5′ UTR and is not tagged at the N-terminus. (B) Western blot analysis of Dicer expression in different tissues of a heterozygote *Dicer*^Δ*HEL1/SOM*^ mouse. (C) Production of endo-siRNAs from MosIR, a dsRNA-expressing plasmid [11] transfected into *Dicer*^ΔHEL1/ΔHEL1^ ESCs. The y-axis depicts RPMs calculated for the entire RNA-sequencing library. (D) Breeding performance of different heterozygous mutants. (E) MA plot of small RNA-seq analysis of whole *Dicer*^*GNT/GNT*^ E15.5 embryos compared to wild type E15.5 embryos. The analysis was performed in biological triplicates. (F) MicroCT scans of E18.5 embryos revealed in *Dicer*^Δ*HEL1/*Δ*HEL1*^ embryos morphological aberrations in heart (right ventricle hypertrophy and loose hypomorphic myocardial walls in both atria), underdeveloped lungs, and growth defect manifested as palate cleft (asterisk, in 4/6 animals), which could also contribute to the perinatal lethality through maternal infanticide of non-feeding pups. (G) MA plot depicting changes in levels of annotated murine miRNAs (miRBase 22.1 [31]). Significantly dysregulated 5p and 3p miRNAs (DESeq p-value 0.05) are shown as oriented blue ▾ and red ▴ triangles, respectively. Mirtrons, as a unique class of non-canonical miRNAs are represented by green triangles whose orientation is the same as that of significantly dysregulated 5p and 3p miRNAs.

To understand the importance of Dicer’s helicase function *in vivo*, we produced mice carrying point mutations in the conserved HEL1 motifs, Walker A (^69^GNT) and Walker B (^175^DQCH), and lacking HEL1 entirely (*Dicer*^Δ*HEL1*^ mutant Fig. 1A and S1). The *Dicer*^Δ*HEL1*^ allele essentially encodes an HA-tagged Dicer^O^ protein. A control allele, designated *Dicer*^*SOM*^, was produced previously [16] to express HA-tagged full-length Dicer while lacking introns 2-6 that were deleted in *Dicer*^Δ*HEL1*^ allele (Fig. 1A and S1H). *Dicer*^Δ*HEL1*^ allele produced the expected truncated Dicer variant (Fig. 1B and S1K) and its functionality was confirmed in embryonic stem cells (ESCs), where it generated ∼10x more siRNAs from long dsRNA than normal Dicer (Fig. 1C).

The catalytically inactive *Dicer*^*GNT/GNT*^ and *Dicer*^*DQCH/DQCH*^ mutant mice were born in the expected Mendelian ratios (Fig. 1D), appeared normal and were fertile. Small RNA analysis of *Dicer*^*GNT/GNT*^ E15.5 embryos revealed only minor changes in the miRNome (Fig. 1E). In contrast, mating of *Dicer*^Δ*HEL1/+*^ animals did not yield any weaned *Dicer*^Δ*HEL1/*Δ*HEL1*^ progeny (Fig. 1D). *Dicer*^*ΔHEL1/ΔHEL1*^ mice died perinatally whereas *Dicer*^*SOM/SOM*^ animals have normal viability [16]. Recovered *Dicer*^*ΔHEL1 /ΔHEL1*^ newborns were cyanotic, had breathing difficulties and had a body weight ∼60% that of heterozygous and wild-type siblings (Fig. S2A-C). Apart from the growth retardation, *Dicer*^*ΔHEL1/ΔHEL1*^ mice had anatomical and physiological aberrations consistent with cardiovascular defects including underdeveloped lungs with reduced branching, heart defects (Fig. 1F), and reduced number of red blood cells and hemoglobin amount per red blood cell (Fig. S2D). In summary, an intact helicase domain rather than its enzymatic activity is required to support Dicer function and postnatal survival.

Detrimental effects of Dicer^ΔHEL1^ protein could be either associated with toxic upregulation of endogenous RNAi or with aberrant miRNA homeostasis. Small RNA analysis of *Dicer*^Δ*HEL1/*Δ*HEL1*^ E15.5 embryos showed strong miRNome dysregulation (Fig. 1G). At the same time, analysis of small RNA clusters in the genome in *Dicer*^Δ*HEL1/*Δ*HEL1*^ E15.5 and *Dicer*^Δ*HEL1/*Δ*HEL1*^ ESCs did not find genomic loci giving rise to abundant pools of siRNAs from long dsRNA (Fig. S3A-B). siRNAs from transcribed inverted repeats in *Optn* and *Anks3* were increased but their abundance in *Dicer*^Δ*HEL1/*Δ*HEL1*^ embryos was still negligible (Fig. S3C). Thus, miRNAs were the most affected abundant class of Dicer-derived small RNAs in the *Dicer*^Δ*HEL1/*Δ*HEL1*^ mutants.

At E15.5, homozygous loss of HEL1 altered the expression of ∼1/4 embryonic miRNAs (386 of 1199 miRNAs with abundance >1 read per million (RPM), Fig. 1G, Table S1) with roughly equal numbers of up-and down-regulated miRNAs. Relative miRNA expression changes correlated well between *Dicer*^Δ*HEL1/*Δ*HEL1*^ embryos and *Dicer*^Δ*HEL1/*Δ*HEL1*^ ESCs (correlation coefficient 0.811, Fig. S3D) suggesting that the miRNome remodeling considerably reflects direct effects of Dicer^ΔHEL1^ on miRNA biogenesis.

Strikingly, a half of the 50 most upregulated miRNAs in *Dicer*^Δ*HEL1/*Δ*HEL1*^ E15.5 embryos were mirtrons, non-canonical miRNAs whose precursors are spliced out specific small introns [17, 18] (Fig. 1G, Table S1). Upregulated mirtron precursors had relatively long stems and/or loops (Fig. 2A), miR-3102 comprising such a long stem that it is carrying two consecutive miRNAs (Fig. 2B-C). The increase in mirtron expression was not transcriptional as mirtron-encoding host genes were not upregulated in *Dicer*^*ΔHEL1/ΔHEL1*^ ESCs (Table S2). Consistent with RNA-seq data, Dicer^ΔHEL1^ cleaved the 5′ radiolabeled pre-miR-7068 (the most upregulated mirtron) *in vitro* more efficiently than normal Dicer (Fig. 2D). Notably, both Dicers cleaved pre-miR-7068 *in vitro* in non-canonical ways, producing also a fragment arising from a single strand cleavage at the 5′ 3p miRNA cleavage site (Fig. 2D). Taken together, Dicer^ΔHEL1^ is more tolerant of extended pre-miRNA stems and loops of mirtrons than the full-length enzyme suggesting that HEL1 physiologically inhibits biogenesis of small RNA from such substrates.

**Figure 2.**
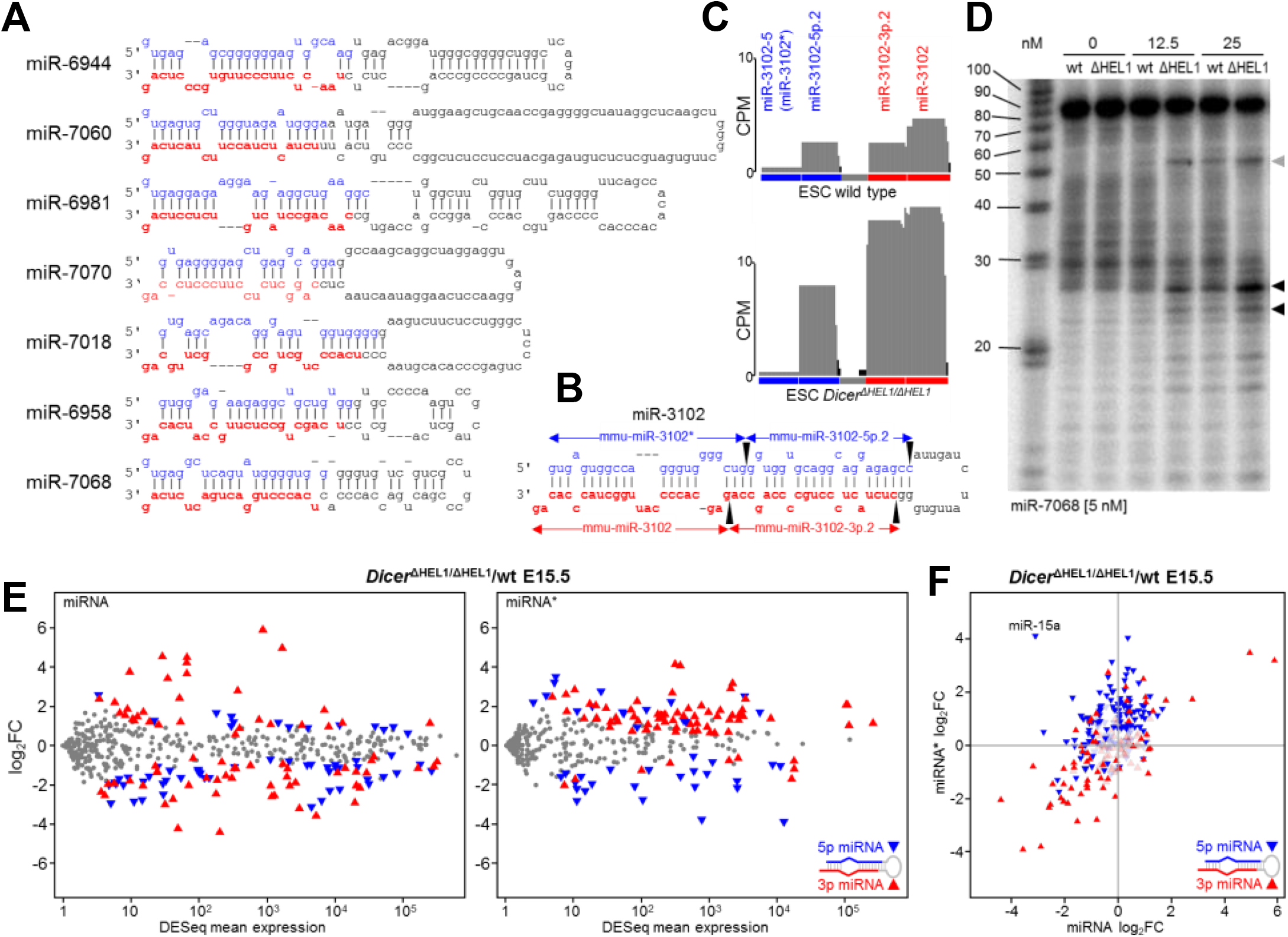
HEL1 restricts processing of mirtrons and 3p passenger strand loading. (A) Strongly upregulated mirtrons have extended stems and larger loops. Secondary structures were adopted from miRBase annotation data. (B) miR-3102 is a unique mirtron, which gives rise to two miRNA:miRNA* duplexes through two consecutive Dicer cleavages at points indicated by black arrowheads. (C) The graphs show changes of miR-3102 levels in *Dicer*^Δ*HEL1/*Δ*HEL1*^ ESCs. The y-scale represents relative expression in counts per million (CPM) estimated by DESeq algorithm. For constructing the graphs, 21-23nt small RNA sequencing data were mapped on the genomic sequence (represented by x-scale as the genomic locus in 5p to 3p orientation), collapsed and normalized per million of 21-23 nt reads. Grey parts represent sequences aligned with the annotated mature miRNA sequence, black parts represent sequences outside the annotated miRNA sequence. (D) *In vitro* cleavage of miR-7068 precursor. Radiolabeled *in vitro* synthesized precursor (final conc. 5 nM) was incubated for 30 min with given concentrations of either full-length Dicer (wt) or with Dicer^ΔHEL1^ (ΔHEL1). Reaction was resolved by PAGE and visualized by phosphorimaging. Black arrowheads depict two cleavage products corresponding to 3′ 5p cleavage positions, a gray arrowhead depicts a product of an asymmetric cleavage. The experiment was repeated three times; a representative gel is shown. (E) MA plots depicting relative changes of dominant miRNAs (left) and passenger strands (miRNA*, right) in *Dicer*^Δ*HEL1/*Δ*HEL1*^ E15.5 embryos. 5p and 3p origins of significantly changed miRNAs or miRNA*s are distinguished by color and triangle orientation as depicted. (F) Relative changes of dominant miRNAs and their passenger strands in *Dicer*^Δ*HEL1/*Δ*HEL1*^ E15.5 embryos. Each triangle depicts the strand (5p or 3p) of the dominant miRNA, its position corresponds to relative changes of the dominant miRNA (x-axis) and its corresponding miRNA* (y-axis). Deep color indicates significantly dysregulated miRNAs.

Another remarkable impact of Dicer^ΔHEL1^ on miRNome concerned passenger strands (miRNA*), the miRNA strands less likely to be loaded onto AGO effector protein after Dicer cleavage. There was striking preferential upregulation of 3p miRNA passenger strands and downregulation of 5p miRNA* in both, *Dicer*^Δ*HEL1/*Δ*HEL1*^ E15.5 embryos (Fig. 2E) and *Dicer*^Δ*HEL1/*Δ*HEL1*^ ESCs (Fig. S3E). A slight reciprocal trend was observed for leading (much more abundant) miRNA counterparts (Fig. 2F and S3F). Since passenger strands have much lower abundance than main strands, their high relative increase would be expected to cause a minor, if experimentally detectable, reduction of corresponding 5p leading miRNAs. However, a strong reciprocal effect resulted in specific cases in a former 3p miRNA* becoming the leading strand, as exemplified by miR-15a and miR-145a (Fig. 3A-B and S4A). Taken together, the loss of HEL1 apparently affects the thermodynamic sensing of the 5′ end of 3p miRNAs and facilitates its selection for AGO loading.

**Figure 3.**
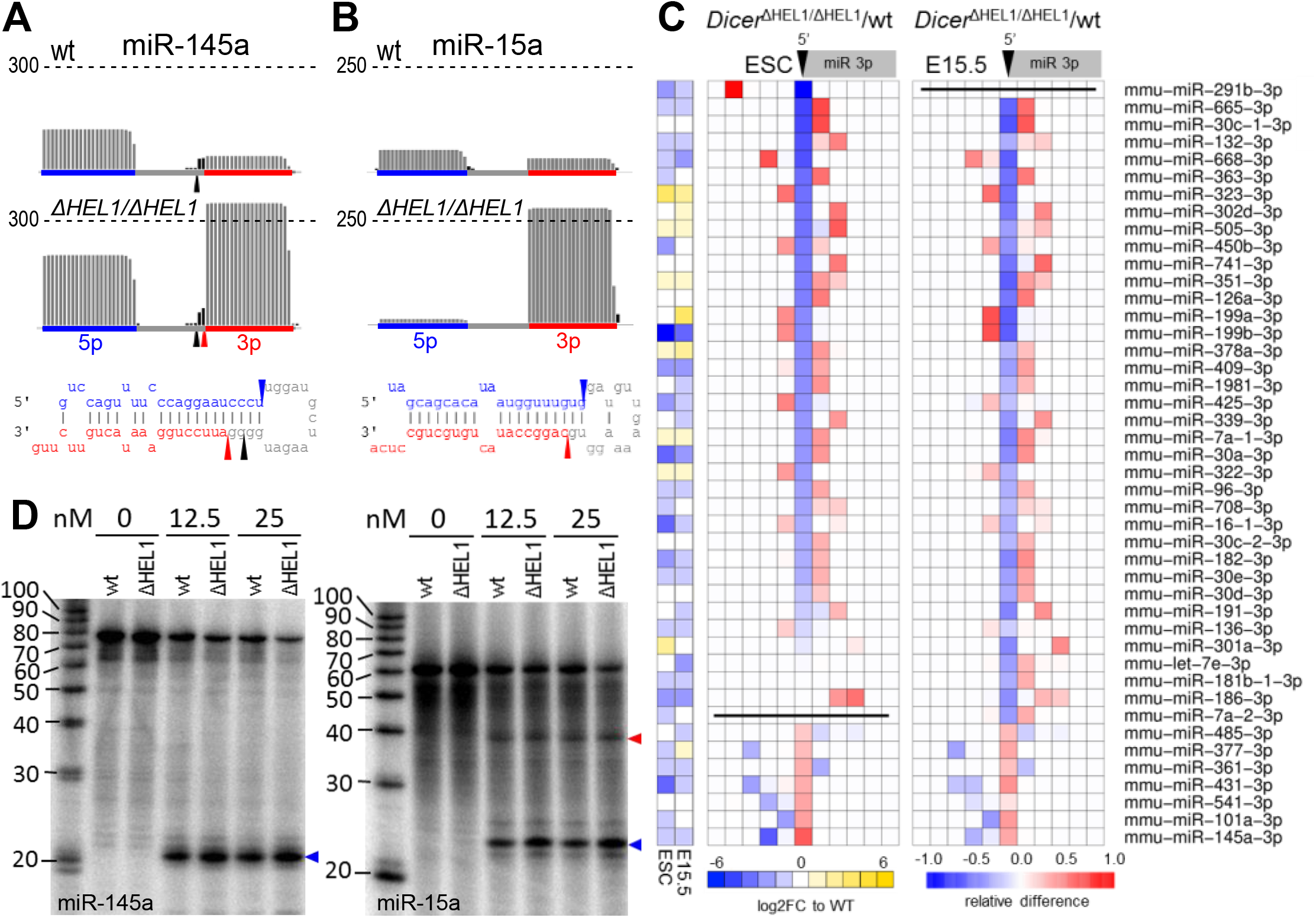
HEL1 is important for pre-miRNA cleavage fidelity. miR-145 (A) and miR-15a (B) exhibit a strong bias towards 3p strand selection in *Dicer*^Δ*HEL1/*Δ*HEL1*^ ESCs. The y-scale represents relative expression in counts per million (CPM) estimated by DESeq algorithm. The graph construction was the same as in Fig. 2C. (C) The heatmap depicts analysis of the 5′ 3p miRNA cleavage site in 50 most affected miRNAs among all 3p miRNAs (>100 DESeq RPMs) in E15.5 embryos and ESCs. The cleavage site is indicated by a black arrowhead. Each column of squares represents one nucleotide from the cleavage site in direction into the mature 3p miRNA (to right) or upstream of it (to left). Red-blue colors indicate relative changes in the 3p miRNA cleavage site when compared with the wild type sample. (D) miR-145a (left panel) and miR-15a (right panel) *in vitro* cleavage assays. 5 nM of *in vitro* synthesized P^32^ 5′-end labeled pre-miRNAs were incubated with indicated concentrations of recombinant Dicer variants at 37°C for 60 minutes, resolved by PAGE, and visualized by phosphorimaging. Blue and red arrowheads point to products corresponding to 5p and 3p cleavage sites in pre-miRNA.

Strand selection was associated with Dicer’s binding partner TARBP2 [19]. Analysis of miRNome in *Tarbp2*^*–/–*^ E15.5 embryos [20] identified 84 differentially regulated miRNAs (>1 RPM, DESeq p-value <0.05, Table S1), majority of which followed a similar trend in *Dicer*^Δ*HEL1/*Δ*HEL1*^ E15.5 embryos (Fig. S4B-C). Since TARBP2 binds the HEL2i subdomain adjacent to HEL1 [21, 22] (Fig. 1a), we examined whether the loss of HEL1 impairs binding of TARBP2 to Dicer^ΔHEL1^. Co-immunoprecipitation of TARBP2 with Dicer showed that TARBP2 remains associated with Dicer^ΔHEL1^ (Fig. S4D) suggesting that miRNome remodeling in *Dicer*^Δ*HEL1/*Δ*HEL1*^ E15.5 embryos is not affected by the loss of interaction between Dicer and TARBP. We hypothesize that HEL1 and TARBP exert similar but non-redundant thermodynamic sensing, which controls 5′ 3p miRNA selection.

Examination of differentially expressed miRNA revealed additional two effects of the loss of HEL1, which can be illustrated on miR-145a and miR-15a, two miRNAs exhibiting a switch of the leading strand (Fig. 3A-B). In case of miR-145a, the strand switch correlated with a shift in the 5′ 3p miRNA cleavage site making the highly abundant miR-145a-3p miRNA in *Dicer*^Δ*HEL1/*Δ*HEL1*^ mutants two nucleotides shorter than the miR-145a-3p miRNA* in wild type cells (Fig. 3A). The loss of two G:C basepairs and presence of 5′ A nucleotide would be expected to favor the miR-145-3p strand selection [23]. This observation prompted a systematic analysis of cleavage fidelity, because changes of the 5′-terminal nucleotides of 3p miRNAs affect nucleotides 2-7, known as the “seed sequence”, which guide target recognition and binding [24]. A change in the seed sequence thus would be biologically significant [25] even if miRNA abundance would not change. Cleavage site shifts at the 5′ end of 3p miRNAs showed a similar pattern in *Dicer*^Δ*HEL1/*Δ*HEL1*^ E15.5 embryos and ESCs (Fig. 3C and S5), suggesting that most of them come from altered cleavage fidelity of Dicer^ΔHEL1^. A 5′ cleavage position change was found in at least 20% of abundant 3p miRNAs in *Dicer*^Δ*HEL1/*Δ*HEL1*^ E15.5 embryos and ESCs and mostly involved a single-nucleotide shift of the cleavage site. Cleavage fidelity was found to be also affected in *Tarbp2*^*–/–*^ embryos [20]. Approximately a half of the cleavage alterations in *Dicer*^Δ*HEL1/*Δ*HEL1*^ samples were also observed in *Tarbp2*^*–/–*^ embryos (Fig. S5) suggesting that the loss of TARBP2 affects biogenesis in a similar but non-redundant mechanism as the loss of HEL1. In contrast, cleavage fidelity in *Dicer*^*GNT/GNT*^ E15.5. mutant was essentially unaffected (Fig. S5). In summary, HEL1 but not its enzymatic activity is important for definition of mature sequences of biologically active miRNAs.

The second effect was increased asymmetric processing of pre-miR-15a, where Dicer^ΔHEL1^ would cleave the pre-miRNA at the 5′ 3p miRNA site while the concurrent cleavage at the 3′ 5p miRNA site would not occur. Partially cleaved miR-15a but not analogous miR-145a fragments were present in RNA-seq data (Fig. S4E). In depth analysis of RNA-seq data identified tens of miRNAs comparable to miR-15a in frequency of RNA fragments from precursors apparently cleaved only at the 5′ 3p miRNA position (Table S3). We confirmed the asymmetric miR-15a precursor cleavage experimentally *in vitro* (Fig. 3D). The partially cleaved miR-15a fragment was also observed upon cleavage with the full-length Dicer (Fig. 3D) suggesting that asymmetric cleavage could be a miRNA-specific feature, which becomes pronounced by Dicer^ΔHEL1^ because of its higher activity and altered thermodynamic sensing. Importantly, miRNA-specific features and diversity of effects on miRNAs in *Dicer*^Δ*HEL1/*Δ*HEL1*^ are not surprising considering stochastic adaptive evolution of individual pre-miRNAs to be bound and processed by the full-length Dicer.

To understand the molecular function of mammalian Dicer’s helicase domain, we used cryo-electron microscopy (cryo-EM) to analyze the two functionally divergent mouse Dicer isoforms (Fig. 4A). We determined the 3.8-Å-resolution structure of mouse full-length Dicer in the apo form (Fig. S6, Table S4). Akin to human Dicer, the overall structure of mouse Dicer shows an identical “L shape” architecture [22], adopting a “closed” state. Cryo-EM data suggest that the helicase domain is flexible around HEL1 consistent with previous observations [22, 26] (Fig. S6). The inherent flexibility of the helicase was greater for Dicer^O^ and prevented determination of its high-resolution structure in the apo form. When we used structural prediction by AlphaFold [27], we obtained a model for Dicer^O^, which showed different relative orientation of the core and the residual helicase domain consisting of HEL2i and HEL2 subdomains (Fig. S7A-C). The helicase domain in Dicer^O^ was rotated by 60° from its position in full-length Dicer, opening up the catalytic center composed of both RNase III cleavage sites (Fig. S7C). This rotation is mainly associated with loss of interactions between the HEL1 subdomain and the RNase IIIb domain. We conclude that HEL1 stabilizes the closed state of Dicer.

**Figure 4.**
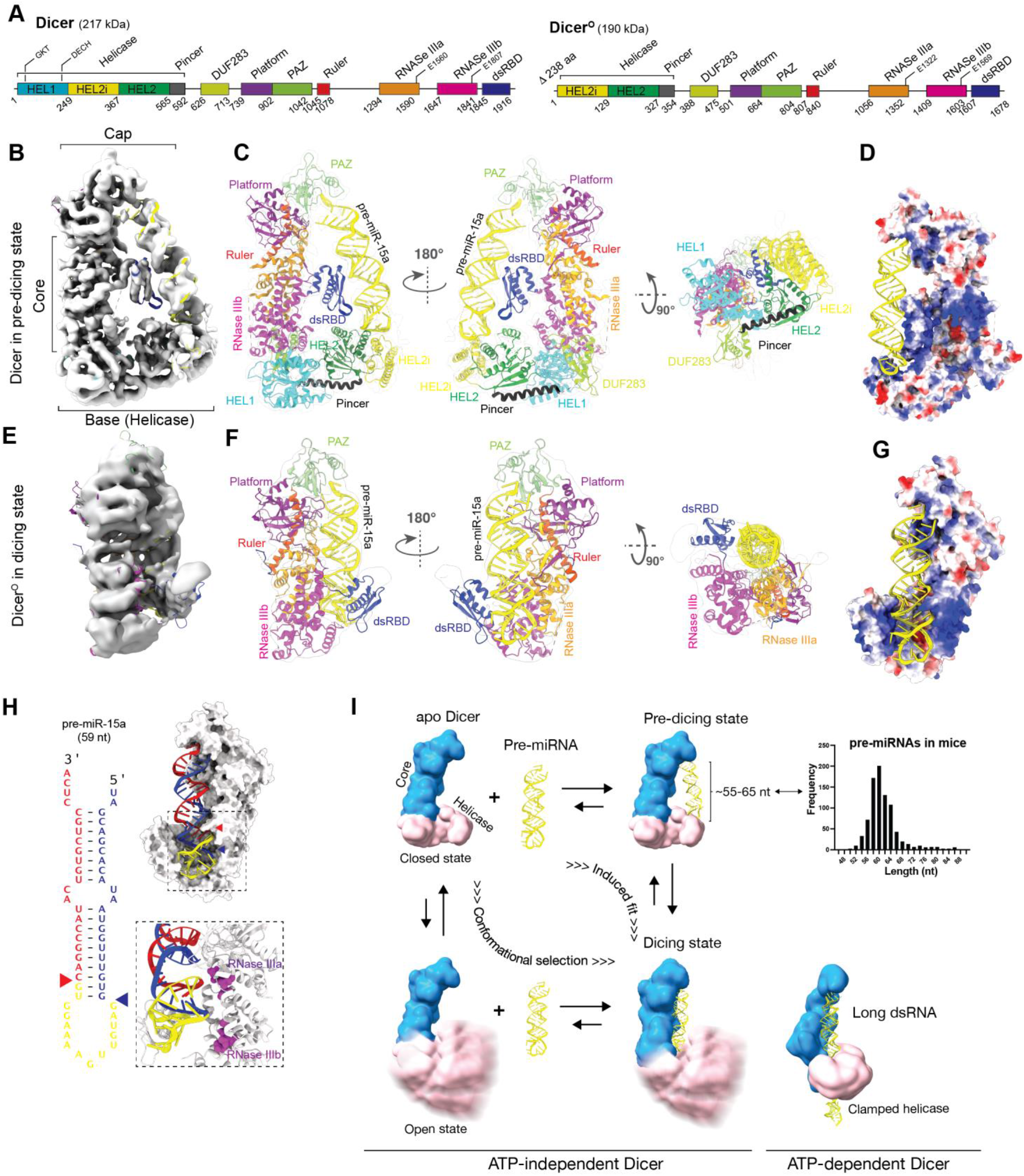
Cryo-EM structures of mouse Dicer and Dicer^O^ reveal the molecular mechanism of how the helicase domain regulates dicing and substrate selection. (A) Domain architecture of Dicer and Dicer^O^ numbered at boundaries. (B-D) Overall structure of the Dicer-RNA complex, shown as 4.19-Å cryo-EM density map (B), in ribbon representations in two orthogonal views (c), and as electrostatic surface view of Dicer showing that the pre-miR-15a (in yellow) is bound by positively charged regions of HEL2, dsRBD, and PAZ-Platform. Blue indicates a positively charged region and red a negatively charged region (D). (E-G) Overall structure of the Dicer^O^- RNA complex, shown as 6.21-Å cryo-EM density map (E), in ribbon representations in two orthogonal views (F), and as electrostatic surface view of Dicer^O^ showing that the pre-miR-15a (in yellow) is bound by positively charged regions of the Dicer^O^ core (G). (H) pre-miR-15a cleavage sites by Dicer^O^. (I) Model of dicing state formation in vertebrate Dicer. Dicer exhibits conformational dynamics at its helicase domain, which exists in two conformations: a major conformation, the closed state; and a minor conformation, the open state. The absence of HEL1 in Dicer^O^ shifts the equilibrium to favor the open state. The dicing state can occur through two pathways: conformational selection or induced fit. In induced fit, Dicer first recognizes the pre-miRNA architecture to form the pre-dicing state and then rearranges the helicase domain to accommodate the pre-miRNA in the dicing state. In conformational selection, Dicer in the open state directly binds the pri-miRNA to form the dicing state, in which the helicase domain remains flexible. All pre-miRNAs are predicted to be accommodated in the Dicer core and are unlikely to provide a sufficient binding platform for the helicase domain. Models are based on experimental data and are shown in a low resolution for clarity. dsRBD is omitted in the models for clarity. (Upper right) Distribution of murine pre-miRNA lengths according to the miRBase 22.1 annotation [31]. (Bottom right) ATP-dependent Dicers mediate processive cleavage by threading of dsRNA through the helicase domain in a clamped conformation (demonstrated by the DCL1 structure (7ELD)).

Next, we determined the 4.19-Å-resolution structure of the complex of the full-length mouse Dicer with a 59-nt precursor of miR-15a (Fig. 4B-D and S8); this miRNA was chosen for its unique behavior in *Dicer*^Δ*HEL1/*Δ*HEL1*^ mutants (Fig. 2F, 3B,D). The PAZ domain anchors the 3′- end of the pre-miR-15a, but not the 5′ end, in contrast to the human enzyme binding pre-let-7 [22, 28], likely reflecting the absence of the human-specific α-helix in the PAZ domain (Fig. S7E). The β-sheet face of the dsRBD binds the central double-helical region of the pre-miR-15a (Fig. 4C), in contrast to other members of the dsRBD family, which typically interact with RNA via their α-helical face [29]. In the Dicer-pre-miR-15a complex, the α-helical face is oriented towards the RNase IIIa/b catalytic sites (Fig. 4C). The large terminal loop of pre-miR-15a binds to the outer rim of the helicase subdomains HEL2i and HEL2 (Fig. 4C-D). These interactions position the pre-miRNA away from the RNase III catalytic sites (Fig. 4C), suggesting that this structure represents a pre-dicing state [22]. The closed state is stabilized by the aforementioned interaction of the HEL1 subdomain and the RNase IIIb domain, which is mediated by aromatic and hydrophobic residues conserved across vertebrates but not in invertebrates (Fig. S7H). The closed state may allow Dicer to recognize specific structural features of miRNA precursors while it would impair processing of mirtrons and dsRNAs because they cannot be optimally recognized due to steric hindrance in the pre-dicing state (Fig. S7F-G).

Since the structural modeling showed that Dicer^O^ adopts the open state, we hypothesized that the structure of Dicer^O^–RNA complex could capture the dicing state. To this end, we determined the 6.2-Å-resolution cryo-EM structure of Dicer^O^ in a complex with pre-miR-15a (Fig. 4E-G and S9, Table S4). The helicase and DUF283 domains showed faint densities and could not be built into the model. A control experiment showed that Dicer^O^ on the grid was intact suggesting that the weak and missing protein densities originated from their inherent flexibility (Fig. S9A). We hypothesize that the helicase domain in ATP-independent Dicers remains flexible in complex with most pre-miRNAs and short dsRNA [30], as they are not expected to protrude far enough from the enzyme core to be clamped analogically to ATP-dependent Dicers from *Arabidopsis* and *Drosophila* (Fig. 4I) [8, 9].

The binding of pre-miR-15a in Dicer^O^ where the dsRBD interacts via its α-helical face, clamping pre-miRNA tightly in the positively charged groove formed by the RNase IIIa/b domains represents the dicing state, which was predicted but never observed in metazoan Dicers (Fig. 4F-G). The dsRBD-RNA contacts span two consecutive minor grooves in the pre-miRNA, while the β1–β2 loop and the N-terminal part of the helix α2 bind to the terminal loop of the pre-miRNA. Our complex structure shows that mouse Dicer^O^ cleaves pre-miR-15a between bases G22 and G23, and between bases G37 and C38, producing a 22-nt miRNA duplex (Fig. 4H), precisely matching pre-miR-15a cleavage sites annotated in the miRBase [31]. Interestingly, the terminal loop of pre-miR-15a interacts with the dsRBD and the RNase IIIb domains but exhibits imperfect alignment with the RNase IIIb catalytic site (Fig. 4H), which is consistent with the asymmetric cleavage of pre-miR-15a at the 5′ end of 3p miRNA described above (Fig. 3D).

In summary, our genetic and structural analyses showed that the intact HEL1 (DExD/H) helicase domain in mouse Dicer has an essential non-canonical function in controlling the equilibrium between Dicer’s closed and open states. We propose a model where the open state, typical for RNAi-supporting Dicer^O^ isoform, can bind a broad range of substrates, including pre-miRNAs, mirtrons, and dsRNAs, and directly form the dicing state (Fig. 4I). The fact that we did not observe Dicer^O^ in a pre-dicing state would explain why Dicer^O^ processes dsRNA more effectively than the full-length enzyme *in vivo* and promotes RNAi [11] (Fig. S10). In contrast, the closed state pre-selects pre-miRNAs and the pre-dicing state may serve as a kinetic trap for diffusion-driven screening of optimal substrates. This gatekeeping function of Dicer in substrate discrimination has profound implications for biogenesis of mammalian small RNAs at the molecular level and, more broadly, for evolution of vertebrate RNA silencing pathways. Highly conserved Dicer’s architecture provides a stable structural “mold” for adaptive evolution of vertebrate pre-miRNAs. This could be a significant factor behind extraordinary expansion of vertebrate miRNA [32], which are stochastically evolving from random RNA structures [33]. At the same time, the gatekeeping function committed mammalian Dicer to its role in the miRNA pathway such that it explains why Dicer^O^ and biologically relevant endogenous RNAi are primarily restricted to mouse oocytes where miRNAs are biologically irrelevant [34].

## Acknowledgments

We thank Vladimir Benes, EMBL sequencing facility and Genomics and Bioinformatics core facility at the Institute of Molecular Genetics for help with RNA-seq experiments, and Kristian Vlahovicek for providing hardware support for bioinformatics analysis. The main funding was provided by the Czech Science Foundation (EXPRO grant 20-03950X to PS and GA22-19896S to RS). Additional funding included the European Research Council (ERC) under the European Union’s Horizon 2020 research and innovation programme (grant agreement No 647403 to P.S. supported developing mouse models and grant agreement No. 649030 to R.S. supported initial experiments). Operational Programme Research, Development and Education-project „Internal Grant Agency of Masaryk University” (No.CZ.02.2.69/0.0/0.0/19_073/0016943 to DZ), and by the institutional support from the Ministry of Education, Youth and Sports (MEYS) of the Czech Republic (CEITEC 2020 project (LQ1601)). Financial support of V.B. and F.H. was in part provided by the Charles University in a form of a PhD student fellowship; this work will be in part used to fulfil requirements for a PhD degree and hence can be considered “school work”. The authors used services of the Czech Centre for Phenogenomics at the Institute of Molecular Genetics supported by the Czech Academy of Sciences RVO 68378050 and by the project LM2018126 Czech Centre for Phenogenomics provided by Ministry of Education, Youth and Sports of the Czech Republic. Authors gratefully acknowledge the cryo-EM Core Facility and Proteomics Core Facility supported by the CIISB research infrastructure, an Instruct-CZ Centre of Instruct-ERIC EU consortium (funded by MEYS CR infrastructure project LM2018127) for their support with obtaining the scientific data presented in this paper. Computational resources for J.P. included support by the project “e-Infrastruktura CZ” (e-INFRA CZ LM2018140) supported by the Ministry of Education, Youth and Sports of the Czech Republic. Computational resources for F.H. included support from the European Structural and Investment Funds grants for the Croatian National Centre of Research Excellence in Personalized Healthcare (contract #KK.01.1.1.01.0010) and Croatian National Centre of Research Excellence for Data Science and Advanced Cooperative Systems (contract #KK.01.1.1.01.0009).

