## Supplementary material for "Molecular basis of indispensable accuracy of mammalian miRNA biogenesis": Material & Methods, Supplementary Tables S2-S6, Supplementary Figures S1-S10

### Supplementary Material & Methods

**Affiliations:**

[1] CEITEC-Central European Institute of Technology, Masaryk University, Brno, Czechia

[2] Institute of Molecular Genetics of the Czech Academy of Sciences, Videnska 1083, 142 20 Prague 4, Czech Republic

[3] Centre for Regenerative Medicine, Institute for Regeneration and Repair, Institute for Stem Cell Research, School of Biological Sciences, University of Edinburgh, 5 Little France Drive, Edinburgh EH16 4UU, UK.

[4] Bioinformatics Group, Division of Molecular Biology, Department of Biology, Faculty of Science, University of Zagreb, 10000 Zagreb, Croatia

[5] Czech Centre of Phenogenomics and Laboratory of Transgenic Models of Diseases, Institute of Molecular Genetics of the Czech Academy of Sciences, v.v.i., Prumyslova 595, 252 50 Vestec, Czech Republic

[6] Wellcome Centre for Cell Biology, School of Biological Sciences, University of Edinburgh, Edinburgh EH9 3BF, UK.

[*] These authors contributed equally

[+] Correspondence to:

### Supplementary Material & Methods.

#### Animals

Animal experiments concerning *Dicer*$^{GNT}$ and *Dicer*$^{DQCH}$ model were carried out in accordance with the Italian law under a license from the Italian Ministry of Health. Animal experiments concerning *Dicer*$^{\Delta HEL1}$ and *Dicer*$^{SOM}$ models were carried out in accordance with the Czech law and were approved by the Institutional Animal Use and Care Committee (approval no. 34-2014).

##### *Genotyping*

Tail biopsies were processed by PCR genotyping kit (Top-Bio) according to the manufacturer's protocol. 1 µl aliquots were used for genotyping PCR using 0.5 U/reaction of DNA polymerase (highQu). Genotyping primers are provided in the Table S6.

##### *Embryo harvest*

Mice were mated overnight, and the presence of a vaginal plug indicated embryonic day (E) 0.5. The embryos were washed in PBS and fixed in 4% PFA.

#### Dicer$^{x}$ mutant mice and ESCs

Production of *Dicer*$^{\Delta HEL1}$ model was analogous to production of *Dicer*$^{SOM}$ (*16*). We first produced ESCs with the *Dicer*$^{\Delta HEL1}$ allele and then used those for producing chimeric mice and establishing *Dicer*$^{\Delta \mathrm{HEL1}}$ line upon germline transmission of the *Dicer*$^{\Delta \mathrm{HEL1}}$ allele. *Dicer*$^{\Delta \mathrm{HEL1}}$ allele in ESCs (*35*) was generated using CRISPR-Cas9 (*36*) mediated modification of the endogenous *Dicer* locus. Pairs of sgRNAs were designed to cleave *Dicer* genomic sequence in intron 2 (sequence of DNA targets: mDcr_i2a 5′-GTACCCAAATGGATAGAA-3′, mDcr_i2b 5′-GTTGGGATGGAGGTTGTT-3′) and intron 6 (sequence of DNA targets: mDcr_i6a 5′-ACTACGCTAGGTGTAAACAG-3′, mDcr_i6b 5′-TGCAGTCCCCGGACGTTAAAT-3′). A template for homologous recombination was designed to contain an HA-tag at the N-terminus of *Dicer* coding sequence fused to exon 7 of Dicer and ~ 1.5 kb overhangs on both ends (Fig. S1A). Final genomic sequence of *Dicer*$^{\Delta HEL1}$ mice is provided in the Supplementary File 1.

To produce *Dicer*$^{\Delta HEL1}$ mouse strain, we first produced mouse chimeras by ESC microinjection into eight-cell – stage embryos (*37*); host embryos were isolated from C57Bl/6NCrl mice (Fig. S1). We used two ESC lines with C57Bl/6NCrl background (commonly used JM8A3.N1 and homemade RS7) and one in 129 strain (R1). For the first three rounds of chimera production, we used homozygous and heterozygous mutant ESCs and obtained mice with varying degree mosaicism, but we failed to obtain transmission of the mutant allele into the next generation. During the fourth round, a heterozygous ESC clone D11 derived from R1 ESC line yielded a male with > 80% chimeric fur. Breeding of this male with ICR females finally lead to germline transmission of *Dicer*$^{\Delta HEL1}$ allele into the next generation and establishment of the *Dicer*$^{\Delta HEL1}$ mouse line. Sequences of the engineered *Dicer* locus in the mouse genome are provided in the

Supplementary File 1. Phenotype analysis was performed with N3 animals, small RNA seq was done with N7 and N8 embryos (all breedings to ICR background).

#### *Dicer$^{GNT}$* mutant mice

The *Dicer$^{GNT}$* allele was generated by replacing wild-type exon 3 with a mutant exon in which Lys60 was mutated to encode asparagine. The *Dicer* locus was targeted with a vector containing homology arms and a *loxP*-flanked neomycin cassette 5′ of exon 3 that contained the Lys60Asn mutation. Southern blotting of genomic SacI-digested DNA from individual ESC-derived clones with a 3′ probe was used to identify homologous recombinants, where the *Dicer$^{GNT\text{-}Neo}$* allele displaying a 5.9-kb DNA fragment could be distinguished from the wild-type allele of 7.1-kb fragment size. Cre-mediated recombination resulted in the excision of the *loxP*-flanked neomycin cassette and the generation of the *Dicer$^{GNT}$* allele. Mice analyzed in this study were on a C57Bl/6 genetic background.

#### *Dicer$^{DQCH}$* mutant mice

The *Dicer$^{FH\text{-}DQCH}$* allele was generated by retargeting the *Dicer$^{Neo}$* allele, which contains a Flag-HA-HA sequence 5′ of exon 2 and a *loxP*-flanked neomycin cassette within intron 2 (*38*). This was achieved with a vector comprised of homology arms, an FRT-flanked hygromycin cassette and exon 5 in which the Glu166 codon was mutated to encode glutamine. Southern blotting of genomic SacI-digested DNA from individual ESC-derived clones with a 3′ probe was used to identify homologous recombinants with the *Dicer$^{FH\text{-}DQCH\text{-}Neo\text{-}Hyg}$* allele displaying a 7.8-kb DNA fragment. Flp-mediated recombination removed the FRT-flanked hygromycin cassette and generated the *Dicer$^{FH\text{-}DQCH\text{-}Neo}$* allele that was identified with the 3′ probe as a 5.9-kb SacI DNA fragment. Cre-mediated recombination led to the excision of the *loxP*-flanked neomycin cassette and the generation of the *Dicer$^{FH\text{-}DQCH}$* allele.

The targeting for both alleles was performed in A9 ES cells. Targeted ES cells were injected into C57BL/6 eight-cell-stage embryos. Targeted mice were crossed to deleter Cre mice (*39*) or FLP-expressing transgenic mice (*40*) to remove antibiotic resistance cassettes. The mice analyzed in this study were on a C57Bl/6 genetic background.

#### Phenotype analyses

##### *Proliferation assay - EdU staining and apoptosis TUNEL assay*

Pregnant mice were injected with 60 μl of 10mM EdU 1.5 hour before embryo harvest at E10.5 and E14.5. The incorporation of EdU was visualized by Click-it EdU Imaging Kit (Invitrogen) in E10.5 whole mount samples and on 7μm paraffin sections from E14.5 embryos. Apoptosis was visualized in whole mount E10.5 embryos by TUNEL method using *In Situ* Cell Death Detection Kit, TMR red (Sigma-Aldrich).

##### *MicroCT*

E18.5 embryos were fixed for 1 week in 4% PFA and stained with Lugol's Iodine solution for 2 weeks. Stock solution (10g KI and 5g I2 in 100ml H2O) was diluted to 25% working solution in water. Stained specimens were embedded in 2.5% low gelling temperature agarose. Scan was performed on SkyScan 1272 high-resolution microCT (Bruker, Belgium), with resolution set to 4 µm.

##### *Hematopoiesis panel*

20 ul of blood from each E18.5 embryo was collected in tube containing anticoagulant EDTA and diluted with 175uL of V-53D Diluent (Mindray, 105-000146-00). The samples were measured in mode Complete blood count with Differentials (CBC + DIFF) on analyzer Mindray 5300 Vet. One-way Anova with Tukey posttest was used for statistical analysis.

#### Cell culture and transfection

Mouse ESCs were cultured in 2i-LIF media: KnockOut-DMEM (ThermoFisher) supplemented with 15% fetal calf serum (Sigma), 1x L-Glutamine (Sigma), 1x non-essential amino acids (ThermoFisher), 50 µM β-Mercaptoethanol (ThermoFisher), 1000 U/mL LIF (Isokine), 1 µM PD0325901, 3 µM CHIR99021 (Selleck Chemicals), penicillin (100 U/mL), and streptomycin (100 µg/mL). All plastic was coated with 1% gelatin (Sigma) in PBS.
NIH 3T3 fibroblasts were cultured in DMEM (Sigma) supplemented with 10% fetal calf serum, penicillin (100 U/mL), and streptomycin (100 µg/mL).
For transfection, cells were plated on 24-well plates, grown to 80% density and transfected using Lipofectamine 3000 (ThermoFisher) according to the manufacturer's protocol. The total amount of transfected DNA was kept constant (1 µg/well).

#### Western blotting

Mouse tissues or ES cells were homogenized mechanically in RIPA lysis buffer supplemented with 2x protease inhibitor cocktail set (Millipore) and loaded with SDS dye. Protein concentration was measured by Bradford assay (Bio-Rad) and 80 µg of total protein was used per lane. Proteins were separated on 5.5% polyacrylamide (PAA) gel and transferred on PVDF membrane (Millipore) using semi-dry blotting for 50 min, 35 V. The membrane was blocked in 5% skim milk in TBS-T, Dicer was detected using anti-HA 3F10 monoclonal primary antibody (High Affinity rat IgG1, Roche #11867431001; dilution 1:500) or anti-HA rabbit primary antibody (Cell Signaling, #3724, dilution 1:1,000) and incubated overnight at 4°C. Secondary anti-Rat antibody (Goat anti-Rat IgG, HRP conjugate, ThermoFisher #31470, dilution 1:50,000) or anti-Rabbit-HRP antibody (Santa-Cruz #sc-2357, dilution 1:50,000) was incubated 1 h at room temperature. For TUBA4A and TARBP2 detection, samples were run on 10% PAA gel and incubated overnight at 4 ºC with anti-Tubulin (Sigma, #T6074, dilution 1:10,000) or anti-TARBP2 (ThermoFisher #LF-MA0209, dilution 1:1,000) mouse primary antibodies. HRP-conjugated anti-mouse IgG binding protein (Santa-Cruz, #sc-525409, dilution 1:50,000) was used for detection. Signal was developed on films (X-ray film Blue, Cole-Parmer #21700-03) using SuperSignal West Femto Chemiluminescent Substrate (Thermo Scientific).

**Immunoprecipitation**

NIH 3T3 cells transfected with plasmids expressing HA-tagged Dicer$^{\Delta HEL1}$ or Dicer$^{SOM}$ variants were lysed in IP Lysis Buffer (10 mM phosphate buffer, pH 7.2, 120 mM NaCl, 1 mM EDTA, 0.5% v/v NP-40, 10% v/v glycerol). Insoluble material was pelleted by centrifugation. Cleared supernatants were diluted 4-times with IP Dilution Buffer (10 mM phosphate buffer, pH 7.2, 100 mM NaCl, 1 mM EDTA, 0.1% v/v NP-40) and incubated with anti-HA magnetic beads (anti-HA mAb, clone #2-2.2.14, ThermoFisher #88836) for 2 h on a rotator. Beads were washed 4-times with IP Dilution Buffer, finally re-suspended in 60 µl water and processed for western blotting. All buffers were supplemented with 1x Protease Inhibitor Cocktail Set (Millipore) and the whole procedure was performed at 4 °C.

**RNA sequencing**

*ESC small RNA-seq*

Cells were plated on 6-well plates and grown to 80 % density. Cells were transfected with 2 µg/well of pCAG-EGFP-MosIR plasmid and cultured for 48 hours. Cells were washed with PBS, homogenized in Qiazol lysis reagent (Qiagen) and total RNA was isolated by Qiazol-chloroform extraction and ethanol precipitation method (*41*). RNA quality was verified by Agilent 2100 Bioanalyzer. Small RNA libraries were constructed using NEBNext Multiplex Small RNA Library Prep Set for Illumina (New England Biolabs) according to the manufacturer's protocol. Small RNA libraries were size selected on 6% PAAGE gel, a band of 140 - 150 bp was cut from the gel and RNA was extracted using Monarch® Genomic DNA Purification Kit. Quality of the libraries was assessed by Agilent 2100 bioanalyzer. Libraries were sequenced on the Illumina HiSeq2000 platform at the Genomics Core Facility at EMBL.

*E15.5 small RNA-seq*

E15.5 embryos were removed from the uterus and washed in PBS. The yolk sac was taken for genotyping and embryos were transferred into RNAlater (Thermo Fisher Scientific). Embryos were homogenized in Qiazol lysis reagent (Qiagen) and total RNA was isolated by Qiazol-chloroform extraction and ethanol precipitation method (*41*). Small RNA libraries were constructed using Nextflex Small RNA-seq kit v3 for Illumina (Perkin Elmer) according to the manufacturer's protocol; 3′ adapter ligation was performed overnight at 20 °C, 15 cycles were used for PCR amplification and NextFlex beads were used for size selection. Final libraries were sequenced by 75-nucleotide single-end reading using the Illumina NextSeq500/550 platform at the core genomics facility of IMG.

#### Bioinformatic analyses

RNA-seq data (Table S5) were deposited in the Gene Expression Omnibus database under accession ID GSE196310 (reviewer access token uhmzccoiprmnbyj)

##### *Mapping of small RNA-seq data*

Small RNA-seq reads were trimmed in two rounds using fastx-toolkit version 0.0.14 (http://hannonlab.cshl.edu/fastx_toolkit) and cutadapt version 1.8.3 (*42*). First, 4 random bases were trimmed from left side:

```
fastx_trimmer -f 5 -i {INP}.fastq -o {TMP}.fastq
```

Next, NEXTflex adapters were trimmed. Additionally, the N-nucleotides on ends of reads were trimmed and reads containing more than 10% of the N-nucleotides were discarded:

```
cutadapt --format="fastq" --front="GTTCAGAGTTCTACAGTCCGACGATCNNNN" --
adapter="NNNNTGGAATTCTCGGGTGCCAAGG" --error-rate=0.075 --times=2 --
overlap=14 --minimum-length=12 --max-n=0.1 --output="$ {TRIMMED}.fastq" --
trim-n --match-read-wildcards $ {TMP}.fastq
```

Trimmed reads were mapped to the mouse (mm10) genome with following parameters:

```
STAR --readFilesIn $ {TRIMMED}.fastq.gz --runThreadN 4 --genomeDir $
{GENOME_INDEX} --genomeLoad LoadAndRemove --readFilesCommand unpigz -c --
readStrand Unstranded --limitBAMsortRAM 20000000000 --outFileNamePrefix $
{FILENAME} --outReadsUnmapped Fastx --outSAMtype BAM SortedByCoordinate --
outFilterMultimapNmax 99999 --outFilterMismatchNoverLmax 0.1 --
outFilterMatchNminOverLread 0.66 --alignSJoverhangMin 999 --
alignSJDBoverhangMin 999
```

##### *miRNA expression analyses*

Mapped reads were counted using program featureCounts (*43*). Only reads with lengths 19-25nt were selected from the small RNA-seq data:

```
featureCounts -a $ {ANNOTATION_FILE} -F $ {FILE} -minOverlap 15 -
fracOverlap 0.00 -s 1 -M -O -fraction -T 8 $ {FILE}.bam
```

The GENCODE gene set (*44*) was used for the annotation of long RNA-seq data. The miRBase 22.1. (*31*) set of miRNAs was used for the annotation of small RNA-seq data. Statistical significance and fold changes in gene expression were computed in R using the DESeq2 package (*45*). Genes were considered to be significantly up- or down-regulated if their corresponding p-adjusted values were smaller than 0.05.

##### *miRNA expression plots – normalization of data, miRNA & miRNA* sorting*

First, the relative position of each mature miRNA ("5p" and "3p" for the miRNA-5p and miRNA-3p, respectively) provided by miRBase 22.1. annotation (*31*) was manually curated and completed. Second, the miRNA type of each mature miRNA ("miRNA" and "miRNA*" for the guide strand and passenger strand miRNA, respectively ) provided by miRBase annotation was completed in this way:

1) The mature miRNAs were assigned into the pair by their hairpin names. The DESeq2 baseMean values of E15.5 and GNT experiments were added to each mature miRNA.
2) The pairs of mature miRNAs with complete miRNA type annotation (both, "miRNA" and "miRNA*" types were present) were preserved.
3) If there is only one mature miRNA annotated in the hairpin, it is assigned as "single_miRNA".
4) If the baseMean values of both miRNAs in the pair are lower than 0.25, it is assigned as "lowExp".
5) For the remaining pairs of mature miRNAs, if the baseMean value of one miRNA is at least double to the second one, it is assigned as "miRNA" / "miRNA*" or "miRNA*" / "miRNA", respectively. Otherwise it is assigned as "notClear".
6) Finally, the newly determined miRNA types are compared to each other. If the mature miRNAs were determined as "miRNA" in one experiment and as "miRNA*" in the other, it is assigned as "cellSpecific". In all the other cases, it there is any discrepancy among the determined miRNA type, it is assigned as "notCLear".

The annotation of the mirtrons was added (*18*).

The DESeq2 baseMean and fold changes were plotted and visualized by home-made R scripts.

The MA plots related to the dominant or passenger strand miRNAs contain only the corresponding miRNAs, all the miRNAs otherwise.

***Small RNA clustering analysis***

Small RNA read clusters (Fig. S3A-B) were identified following the algorithm used in previous studies (*11, 46*). Briefly:

1) Reads were weighted to fractional counts of 1/n where n represents the number of loci to which read maps
2) Reads were then collapsed into a unified set of regions and their fractional counts were summed
3) Clusters with less than 3 reads per million (RPM) were discarded
4) Clusters within 50 bp distance of each other were joined

Only clusters appearing in all replicates of the same genotype (intersect) were considered in the final set. Union of coordinates of overlapping clusters were used to merge the clusters between the samples. Clusters were then annotated, and if a cluster overlapped more than one functional category, the following classification hierarchy was used: miRNA > transposable elements > mRNA (protein coding genes) > misc. RNA (other RNA annotated in ENSEMBL or repeatMasker) > other (all remaining annotated or not annotated regions).

***Cleavage fidelity analysis***

Only miRNAs with DESeq2 baseMean values >= 100 were selected. The cleavage points' coordinates (CP) were extracted from their miRBase 22.1 annotation (*31*). The reads of the lengths 19-25nt were selected from each replicate library. The starting and ending position of all reads were summed up in the CP and its vicinity (+/-15nt) and assigned as 3′-CP of miRNA-5p and 5′-CP of miRNA-3p, respectively. Then, the canonical miRBase CPs were re-defined based on our wild-type data:

1) Position with maximal counts (median among replicates) is assigned as the new CP.
2) If the new CP is more than 7nt outside the canonical one, keep the canonical one.
3) If there are multiple CPs with the same max counts, keep the canonical one.
4) If there are no data / no reads, keep the canonical one.

The counts were extracted for each miRNA at the position of the newly defined CP with 5nt flanks on each side. The read counts were re-calculated into read densities. The final matrix was achieved as a subtraction between a mutant and its corresponding wild-type control. Top 50 miRNAs from *Dicer* mutants were selected based on the absolute value of the difference at the position of CP. Selected miRNAs were ordered by the change of *ESC* fidelity at the position of CP.

***Partial processing analysis***

All sequence reads were selected that overlapped the corresponding pre-miRNA locus in the sense direction. All coordinates (starting/ending position of the miRNA-5p/-3p) were extracted from the miRBase 22.1 annotation (*31*). The categories shown in the Fig.5D and S5D were defined by pre-miRNA boundaries and the two annotated Dicer cleavage points (deviation of the boundaries +/-2nt allowed). Each read was unambiguously assigned into the appropriate category. The percentage from the total number of overlapping reads was calculated.

#### Recombinant plasmid preparation

pCIneo plasmid carrying human DICER1 (NM_1777438) was prepared by standard molecular cloning procedures. The C-terminal 2× FLAG tag and deletion (dHEL1, dHEL2 and dDExD) variants were prepared using Q5 Site-Directed Mutagenesis Kit (NEB) according to the manufacturer’s instructions.

pFastBac plasmids carrying recombinant mouse full-length Dicer and short variant ($Dicer^{O}$) were prepared as follows. The N-terminal fragment containing TwinStrep and HA tags together with TEV protease cleavage site was PCR amplified and inserted into BamHI-SalI restriction sites in pFastBACT1 plasmid (Invitrogen). Subsequently, the C-terminal fragment containing 2xFLAG and 8xHis tags together with TEV protease cleavage site was PCR amplified and inserted into NotI-HindIII restriction sites.

Mouse Dicer and $Dicer^{O}$ omitting start and stop codons were PCR-amplified from pEF1-MH.Bl-$mDcr^{SOM}$ (Addgene) and pEF1-MH.Bl-$mDcr^{OO}$ (Addgene) plasmids, respectively, and inserted in-frame into SalI-NotI sites of the modified pFastBACT1 plasmid using common cloning techniques. C-terminal 2× FLAG tag and deletion variants were prepared using Q5 Site-Directed Mutagenesis Kit (NEB) according to the manufacturer’s instructions (PCR primers: Twin-HA-

TEV_Fwd, Twin-HA-TEV_Rev, 3C-FLAG-His_Fwd, 3C-FLAG-His_Rev, mDicer_SalI_Fwd, mDicerO_SalI_Fwd, mDicer-NotI_Rev).

The catalytically inactive variants of Dicer/Dicer$^{O}$ were prepared by mutating the key residues E1560 and E1807 of the RNAseIII domains into alanine residues (Zhang et al., 2004b) using Q5 Site-Directed Mutagenesis Kit (NEB) kit according to the manufacturer's instructions (PCR primers: mDicer E1560A Forward, mDicer E1560A Reverse, mDicer E1807A Forward, mDicer E1807A reverse). List of all used oligonucleotides can be found in Table S3. All constructs were verified by sequencing.

#### Western blotting

U-2 OS cells transfected with DICER1 variants were lyzed in RIPA lysis buffer supplemented with 1× protease inhibitor cocktail set (Millipore) and loaded with SDS dye. Protein concentration was measured by Bradford assay (Bio-Rad) and 100 µg of total protein was used per lane. Proteins were separated on a 6% SDS-PAGE gel and transferred onto a PVDF membrane (Millipore/Merck) using semi-dry blotting. The membrane was blocked with 5% skimmed milk in TTBS. The DICER1 variants were detected using anti-FLAG mouse monoclonal M2 antibody (Sigma, dilution 1:10,000) and incubated overnight at 4 ºC. HRP-conjugated anti-mouse IgG binding protein (Santa-Cruz, dilution 1:50,000) was used for detection of the primary antibody. Signal was developed on films (X-ray film Blue, Cole-Parmer) using SuperSignal West Femto Chemiluminescent Substrate (ThermoFisher Scientific).

#### Luciferase assay

Dual luciferase activity was measured according to Hampf and Gossen (*47*) with some modifications. Briefly, cells were washed with PBS and lyzed in PPTB lysis buffer (0.2% v/v Triton X-100 in 100 mM potassium phosphate buffer, pH 7.8). A 3-5 µl aliquots were used for measurement in 96-well plates using Modulus Microplate Multimode Reader (Turner Biosystems). First, firefly luciferase activity was measured by adding 50 µl substrate (20 mM Tricine, 1.07 mM $(MgCO_3)_4 \cdot Mg(OH)_2$, 2.67 mM $MgSO_4$, 0.1 mM EDTA, 33.3 mM DTT, 0.27 mM Coenzyme A, 0.53 mM ATP, 0.47 mM D-Luciferin, pH 7.8) and signal was integrated for 10 sec after a 2 sec delay. Signal was quenched by adding 50 µl *Renilla* substrate (25 mM $Na_4PP_i$, 10 mM Na-Acetate, 15 mM EDTA, 500 mM $Na_2SO_4$, 500 mM NaCl, 1.3 mM $NaN_3$, 4 µM Coelenterazine, pH to 5.0) and *Renilla* luciferase activity was measured for 10 sec after a 2 sec delay. Hairpin-expressing plasmids and luciferase reporters are described and deposited in Addgene. RlucIR plasmid expressing a hairpin structure targeted to *Renilla* luciferase coding region was prepared similarly to MosIR using common cloning techniques.

#### Protein purification

The coding sequence and the necessary regulatory sequences of mouse Dicer variants were transposed into bacmid using *E. coli* strain DH10bac. The viral particles were obtained by transfection of the bacmids into the *Sf*9 cells using FuGENE Transfection Reagent (Promega) and further amplification in *Sf*9 cells. Dicer variants were expressed in 200 ml of Hi5 cells (infected at $1.2\times10^6$ cells/ml) with the corresponding P1 virus at multiplicity of infection >1. The cells were harvested 48 hours post infection, washed by 1x PBS, and stored at -80°C. Subsequent operations were carried out at + 4ºC. Pellets were resuspended in ice-cold lysis buffer containing 50 mM Tris (pH 8.0), 300 mM NaCl, 0.4% Triton X-100, 10% (v/v) glycerol, 10 mM imidazole, 1 mM DTT, 2 mM $MgCl_2$, benzonase (250U), and protease inhibitors (0.66 µg/ml pepstatin, 5 µg/ml benzamidine, 4.75 µg/ml leupeptin, 2 µg/ml aprotinin) (Applichem). The resuspended cells were gently shaken for 10 min at 4°C. To aid the lysis, cells were briefly sonicated. The lysate was cleared by centrifugation at 21,000xg for 1 hr at 4ºC. The supernatant was passed through a column containing 2.5 ml NiNTA-agarose (QIAGEN). The affinity matrix was washed 5-times with 15 ml of washing buffer (50 mM Tris (pH 8.0), 500 mM NaCl, 1 mM DTT, 2 mM $MgCl_2$, and 10 mM imidazole). The protein was eluted three times with 3.5 ml of elution buffer (50 mM Tris (pH 8.0), 500 mM NaCl, 1 mM DTT, 2 mM $MgCl_2$, and 300 mM imidazole). The fractions containing protein were pooled and concentrated to 1 ml using 100 kDa cut-off Vivaspin Turbo15 (Sartorius). The proteins were further purified on a size exclusion column (Superose 6 Increase 10/300 GL, GE Healthcare) equilibrated with a buffer containing 50 mM Tris (pH 8.0), 150 mM NaCl, 1 mM DTT, 2 mM $MgCl_2$. Fractions containing protein were pooled, concentrated, snap-frozen in liquid nitrogen, and stored at -80ºC until further use.

#### Radiolabeling of RNA

The RNA was diluted to 250 nM with nuclease-free water and mixed with T4 Polynucleotide Kinase buffer. The RNA was refolded by heating the mixture at 95°C for 3 min and snap-cooled on ice for 5 min. After addition of RNAse inhibitors (NEB), T4 polynucleotide kinase (NEB), and [$\gamma$-$^{32}$P]-ATP (HARTMANN ANALYTIC), the reaction was incubated at 37°C for 10 minutes. The 5´-radiolabeled RNA was purified on G-25 columns (Cytiva) and diluted to a final concentration of 50 nM. The radiolabeled RNA Decade Marker (ThermoFisher Scientific) was prepared according to the manual. The RNA and the marker were aliquoted and stored at -20°C.

#### *In vitro* cleavage assay

*Substrate preparation:* In vitro synthesized RNA oligonucleotides were diluted to 250 nM with nuclease-free water and mixed with T4 Polynucleotide Kinase buffer. The RNA was refolded by heating the mixture at 95°C for 3 min and snap-cooled on ice for 5 min. After addition of RNase inhibitors (NEB), T4 polynucleotide kinase (NEB), and [$\gamma$-$^{32}$P]-ATP (HARTMANN ANALYTIC), the reaction was incubated at 37°C for 10 minutes. The 5′-radiolabelled RNA was purified on G-25 columns (GE Healthcare) and diluted to a final concentration of 50 nM. The radiolabelled RNA Decade Marker (ThermoFisher Scientific) was prepared according to the manual. The RNA and the marker were aliquoted and stored at -20°C.

*Nuclease-activity assay:* Time-course experiments were performed in 10 µl, containing 5 nM labelled RNA substrate, and 100 nM Dicer$^{SOM}$ and Dicer$^{\Delta HEL}$, respectively, in 30 mM Tris (pH

7.0), 30 mM NaCl, 1 mM DTT, and 2 mM $MgCl_2$ at 37ºC. Increasing concentrations (12.5, 25, and 50) of Dicer$^{SOM}$ and Dicer$^{\Delta HEL1}$, respectively, were mixed with 5 nM labelled RNA substrate in 30 mM Tris (pH 7.0), 30 mM NaCl, 1 mM DTT, and 2 mM $MgCl_2$. After 60 min incubation at 37ºC, the reactions were stopped with equal volume of 95% formamide, boiled for 5 min, and analyzed on a 20% polyacrylamide gel containing 8 M urea.

After electrophoresis, the gels were exposed for 6-18 hours onto a phosphor imaging screen (Fujifilm). The signal was detected using FLA 9000 phosphorimager (Fujifilm) and analyzed in Multi Gauge v3.2 software.

##### ***In vitro*** **reconstitution of the Dicer–pre-miR-15a complex**

To refold pre-miR-15a RNA, it was heated for 3 min at 95°C and snap-cooled on ice for 5 min. The complex was formed by mixing 1.5 nmol of pre-miR-15a and 0.5 nmol of catalytically inactive Dicer or Dicer$^{O}$ variant in 50 µl of 50 mM Tris (pH 8.0), 100 mM NaCl, 1 mM DTT, and 2 mM $MgCl_2$. After 30 min incubation on ice, the mixture was applied onto Superose 6 Increase 5/150 GL (Cytiva) column attached to an ÄKTA Purifier (Cytiva). Fractions containing the complex were collected and concentrated to 0.2 mg/ml. The purity and homogeneity of the protein was assessed by SDS-PAGE, while RNA was verified by denaturing gel electrophoresis (20% polyacrylamide gel containing 8 M urea) and visualized using SYBR Gold dye (ThermoFisher Scientific).

##### **Cryo-EM specimen preparation and data acquisition**

The purified Dicer or Dicer–pre-miR-15a complex were diluted to a concentration of about 1 µM in a buffer containing 50 mM Tris (pH 8.0), 100 mM NaCl, 1 mM DTT, and 2 mM $MgCl_2$. The Lacey carbon M300 grid (SPI supplies) was glow-discharged (15 sec, hydrogen-oxygen) immediately before preparing the cryo-EM specimen. In a Vitrobot Mark IV (ThermoFisher Scientific), 3.5 µl of the protein–RNA complex was applied on the grid from the plasma treated side. The grid was blotted for 5.0 sec, blot force -3, in 100% humidity at 4°C, and plunged in liquid ethane cooled by liquid nitrogen. For Dicer, UltraAuFoil M300 (R1.2/1.3) grid (Quantifoil) was glow-discharged (60 sec, argon-oxygen) and 3.5 µl of the protein was applied from the plasma treated side. The grid was blotted for 3.0 sec, blot force 0 in 100% humidity at 4°C. The data were collected using Titan Krios (ThermoFirsher Scientific) transmission electron microscope using SerialEM software (Mastronarde, 2005). The details about data acquisition, processing, structural refinement and validation are shown in Table 4.

##### **Image processing of electron micrographs**

The movies were first processed by MotionCor2 (*48*) for generation of motion corrected, dose-weighted micrographs stack. The CTF parameters were estimated using GCTF (*49*). The micrographs were further manually curated to select for Astigmatism lower than 800 Å and CTF fit parameter lower than 4.5 Å. For each dataset, a set of 30-50 randomly selected micrographs was used for manual particle picking using e2boxer.py tool from EMAN2 (*50*) package. The manually picked particles were used for model generation using crYOLO (*51*). The particles obtained from full dataset picking were imported into cryoSPARC (*52*). Further analysis comprised following steps, 2D classification, *ab-initio* modelling and 3D Refinement. The initial volume maps were further used as a reference for re-analysis of the data using 3D Classification in Relion 3.1 (*53*) and/or training of TOPAZ (*54*) tool to improve the quality of particle picking procedure. The final 3D Refinement was performed in cryoSPARC software. The final maps were further processed using DeepEMhancer (*55*). To further improve the resolution of the Dicer cryoEM map, HEL1+HEL2i+HEL2 domains (Dicer core masked), and the Dicer core (HEL2i+HEL2 masked) were individually locally refined. These locally refined maps were processed using DeepEMhancer and combined for subsequent model building. The detailed statistics are available in Table 4.

**Cryo-EM model building and refinement**

Initial PDB coordinates of the Dicer structure were taken from AlphaFold database (*27*). Regions of low confidence prediction (pLDDT < 50) were excluded from the structure and the remaining blocks of the coordinates were fitted into Dicer sharpened electron density map obtained from DeepEMhancer (*55*) using UCSF Chimera's tool 'Fit in Map' (*56*). The PDB coordinates and the electron density map were then imported into program Coot (*57*) and the tool 'Real Space Refine Zone' was used to achieve optimal fit of the PDB coordinates within the map. Low resolution regions and regions where the map was lacking electron density were excluded from the structure. The dsRBD was docked into map with rigid body approach and fit was optimized using Phenix 'rigid_body' strategy (*58*). The coordinates were validated using Coot's tools 'Ramachandran Plot', 'Rotamer Analysis', and 'Density Analysis'. The same procedure was applied to Dicer–pre-miR-15a complex sharpened electron density map from DeepEMhancer. The initial coordinates of pre-miR-15a were obtained from a modeling server RNAComposer (*59, 60*). The model was fitted and refined into the electron density map using ProSMART Self Restraints implemented in Coot software. The optimized coordinates of the Dicer structure and the Dicer–pre-miR-15a complex were individually subjected to further structural refinement with Phenix software and ISOLDE (*61*). To maintain optimal RNA geometry, stacking and hydrogen bonds restraints were provided in Phenix's 'Real-space refinement'. Initial PDB coordinates of the Dicer$^{O}$ structure were predicted by AlphaFold software. After excluding low confidence prediction regions (pLDDT < 50), the structure was fitted into Dicer$^{O}$–pre-miR-15a complex sharpened electron density map obtained from CryoSparc as described above. Protein domains that were not resolved within the electron density map (residues 1-500) were excluded from the model. Modelled pre-miR-15a was manually fitted into electron density map. ISOLDE's bulk flexible fitting approach with distance restrains was applied after the structure was divided into separate blocks (namely consisting of dsRBD, pre-miR-15 and the rest of the Dicer$^{O}$ structure). MolProbity and PDB Validation tool was used to obtain the overall refinement and structural statistics.

**Quantification and Statistical Analysis**

For the quantification of the EMSA assays, the analyses were carried out using the Multi Gauge v3.2 software (Fujifilm). GraphPad Prism was used to plot the obtained values (Specific binding with Hill slope) and perform the statistical analysis. The bound fraction was determined as the disappearance of the signal corresponding to the unbound substrate (Lane 0). Each data point represents an average of at least two independent experiments. Error bars represent standard deviation (SD).

**Data visualization**

Molecular graphics images were produced using the UCSF Chimera (*56*) and ChimeraX (*62*) package from the Resource for Biocomputing, Visualization, and Informatics at the University of California, San Francisco (supported by NIH P41 RR-01081) and/or Coot (*57*).

**Data availability**

RNA sequencing data were deposited to Gene Expression Omnibus (GEO) with the following accession numbers GSE196310.

All original codes have been deposited at https://github.com/fhorvat/bioinfo_repo/tree/master/papers/2022.DicerX_invivo_paper and are publicly available as of the date of publication.

Any additional information required to reanalyze the data reported in this paper is available from the lead contact upon request.

The accession numbers of the 3.84 Å resolution EM map of mouse Dicer, the 4.19 Å resolution EM map of mouse Dicer–pre-miR-15a complex and the 6.21 Å resolution EM map of mouse Dicer$^{O}$–pre-miR-15a complex and their corresponding coordinates reported in this paper are EMDB: 14387, 14383, and 14384, and PDB: 7YZ4, 7YYM, 7YYN, respectively.

#### Supplementary Figures

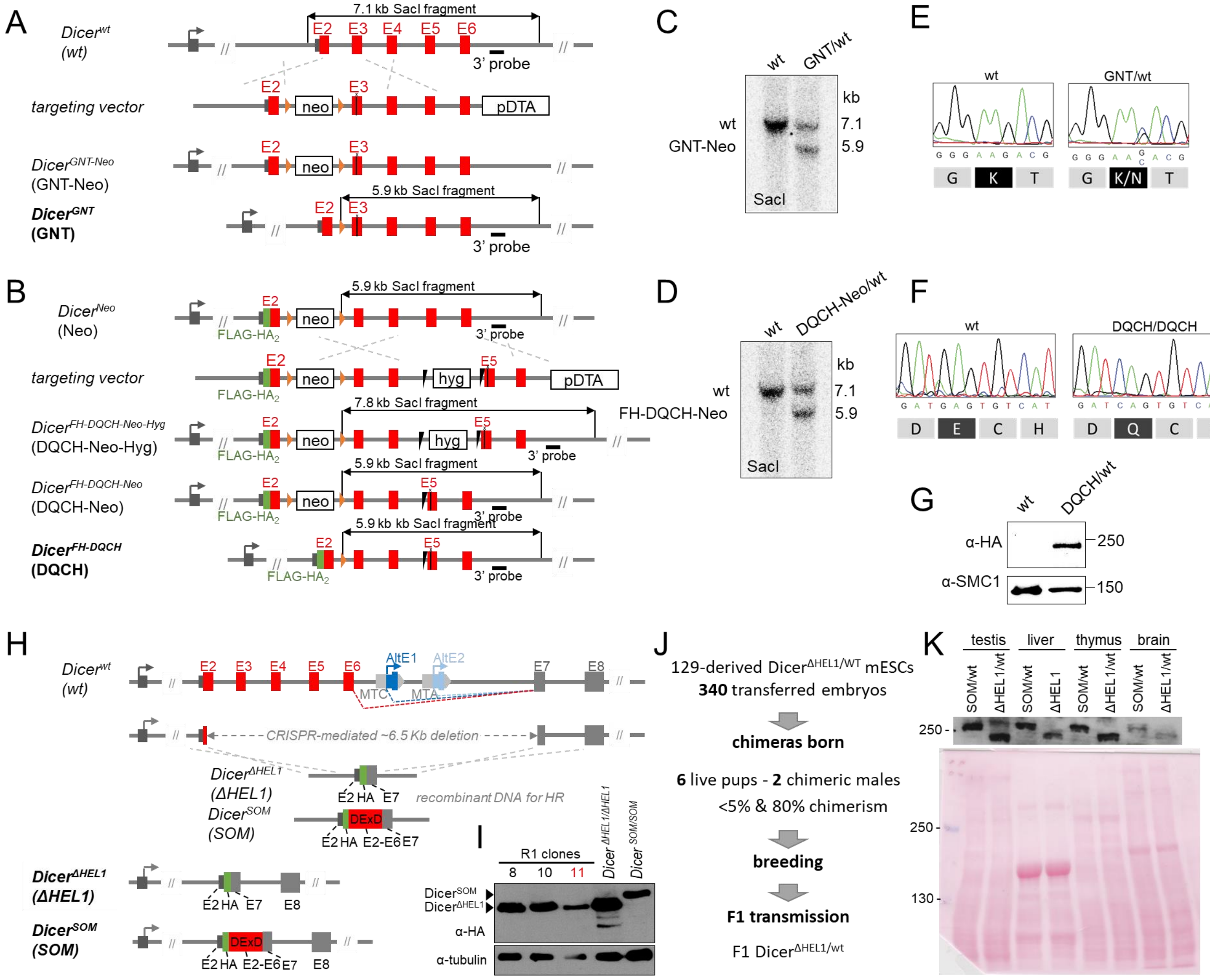


**Fig. S1. Production and validation of Dicer mouse mutants.**

(**A, B**) Schematic depiction of introduction of the GNT and the DQCH mutation into endogenous *Dicer* gene, respectively. (**C**) Detection of the GNT and (**D**) DQCH alleles by Southern blotting. (**E, F**) Validation of the mutated alleles by Sanger sequencing. (**G**) Western blot demonstrating expression of the mutated Dicer$^{DQCH}$ protein. (**H**) Schematic depiction of engineering of *Dicer*$^{\Delta HEL1}$ (ΔHEL1) and *Dicer*$^{SOM}$ (SOM) in the genomic sequence encoding HEL1 of the endogenous *Dicer* gene. Briefly, a fragment from exon 2 to exon 7 was removed using CRISPR /Cas9 and recombined with a *Dicer*$^{\Delta HEL1}$ recombination construct carrying exon 2 (5′ UTR and start codon), HA-tag, and exon 7 coding sequence. *Dicer*$^{SOM}$, which was produced using the same strategy, was described previously(*16*). (**I**) Western blotting of selected positive clones using anti-HA antibody, tubulin was used as a loading control. The heterozygous line 11 gave rise to *Dicer*$^{\Delta HEL1}$ mice. (**J**) Outline of the *Dicer*$^{\Delta HEL1}$ mouse strain production process. (**K**) Western blot analysis of Dicer$^{\Delta HEL1}$ expression in different tissues of a heterozygote Dicer$^{\Delta HEL1/wt}$ mouse. Tissues from Dicer$^{SOM/wt}$ mouse were used for comparison. 80 µg of total protein lysate were loaded per lane. Ponceau staining of the membrane shown below provides control for equal loading.

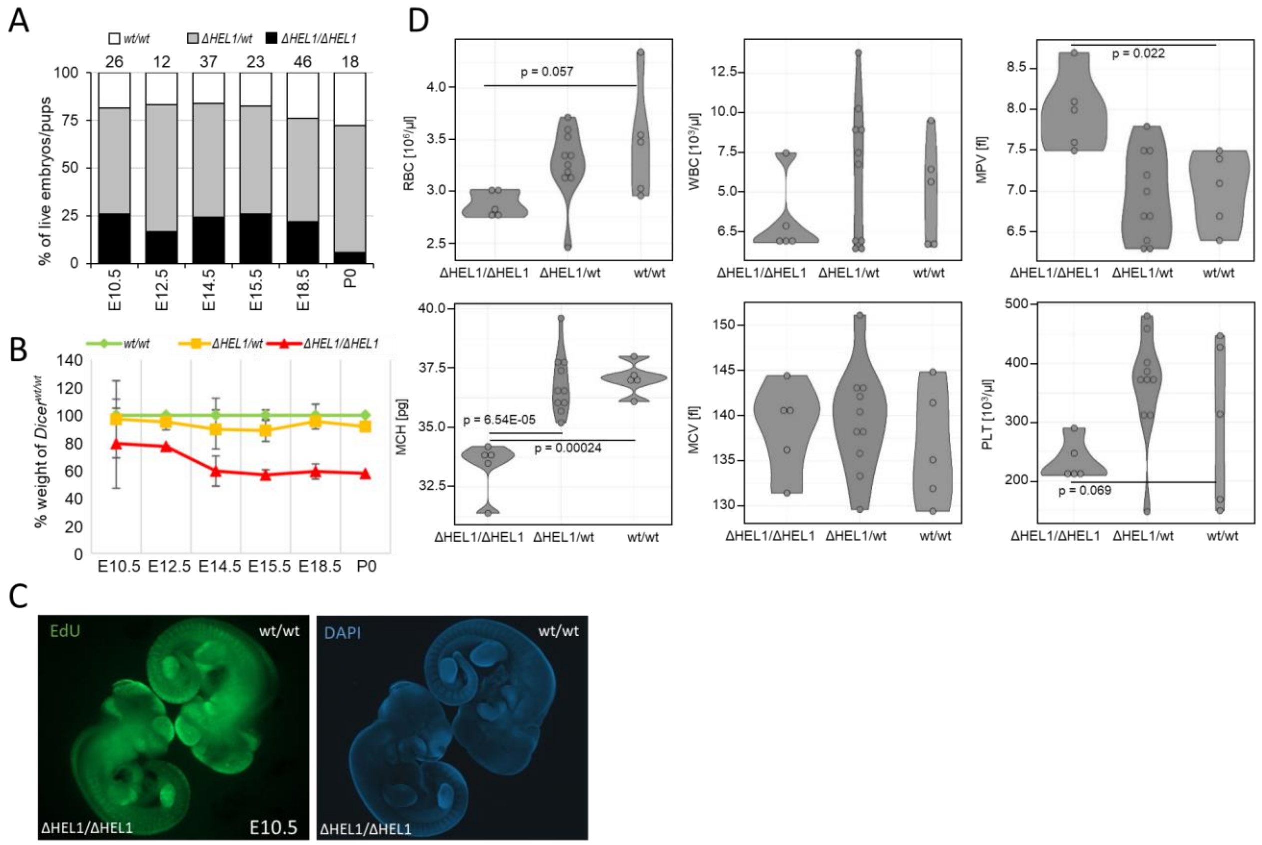


**Fig. S2. Main phenotype characterization of *Dicer*$^{ΔHEL1/ΔHEL1}$ mouse mutants.**

(**A**) Analysis of genotype segregation in embryos from *Dicer*$^{ΔHEL1/wt}$ parents. *Dicer*$^{ΔHEL1/ΔHEL1}$ embryos survive until birth but die soon after. (**B**) Weight of embryos normalized to wild-type shows relative retardation of *Dicer*$^{ΔHEL1/ΔHEL1}$ embryos – the observed growth retardation does not appear to be a simple proliferation defect. While a lower weight in *Dicer*$^{ΔHEL1/ΔHEL1}$ embryos becomes apparent at E10.5, the main reduction of growth appears between stages E12.5 and E14.5, but later it becomes comparable to wild-type controls. (**C**) Analysis of cell proliferation in E10.5 embryos by a combined EdU and DAPI staining shows no apparent difference in general proliferation pattern suggesting more specific mechanism beyond the growth retardation phenotype. The experiment was performed twice with the same result. (**D**) Defects in hematopoiesis found in *Dicer*$^{ΔHEL1/ΔHEL1}$ mice. Shown are: red blood cell count per µl (RBC, reduced by 17%), mean red blood cell volume (MCV), mean hemoglobin per cell (MCH, reduced by 10%), platelet count per µl (PLT), mean platelet volume (MPV), and white blood cell count (WBC) per µl.

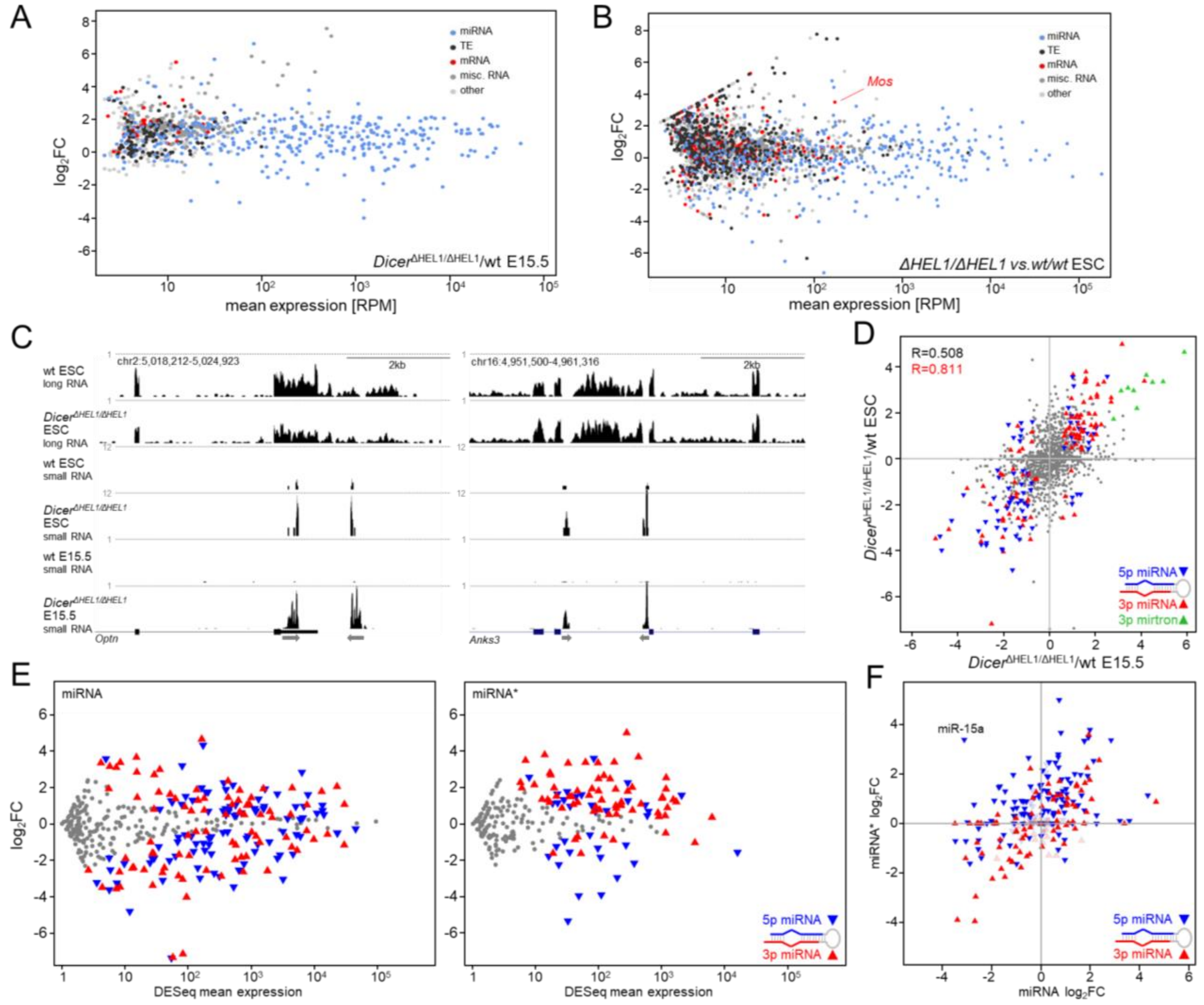


**Fig. S3. miRNome dysregulation in $Dicer^{\Delta HEL1/\Delta HEL1}$ mouse mutants.**

Effect of $Dicer^{\Delta HEL1}$ expression on small RNAs in E15.5 embryos (**A**) and in ESCs (**B**). Each MA plot shows results of small RNA-seq analysis of a $Dicer^{\Delta HEL1/\Delta HEL1}$ sample compared with the normal control (wild-type siblings or the parental ESC line). Each colored point represents a genomic region (cluster) producing 21-23 nt RNAs. Clusters were identified and categorized by a previously developed algorithm(*11*). Cluster "expression" is defined as a fraction of small RNA reads mapping to it per million 21-23 nt small RNAs (RPM). (**C**) UCSC browser snapshots of small RNA populations in *Optn* and *Anks3* loci, which contain transcribed inverted repeats (indicated by gray arrows). Maximum values shown above tracks are counts per million (CPM). (**D**) Relative changes of significantly differentially expressed miRNAs in $Dicer^{\Delta HEL1/\Delta HEL1}$ E15.5 embryos (shown as colored triangles, other miRNAs are depicted as grey circles) correlate with changes of these miRNAs in $Dicer^{\Delta HEL1/\Delta HEL1}$ ESCs. Axes depict $log_2$FC. (**E**) Analysis of miRNA and passenger strand changes in $Dicer^{\Delta HEL1/\Delta HEL1}$ ESCs. MA plots depict relative changes of dominant miRNAs (left) and passenger strands (miRNA*, right) in $Dicer^{\Delta HEL1/\Delta HEL1}$ ESCs. 5p and 3p origins of significantly changed miRNAs or miRNA*s are distinguished by color and triangle orientation as depicted. (**F**) Relative changes of dominant miRNAs and their passenger strands in $Dicer^{\Delta HEL1/\Delta HEL1}$ ESCs. Each triangle depicts the strand (5p or 3p) of the dominant miRNA, its position corresponds to relative changes of the dominant miRNA (x-axis) and its corresponding miRNA* (y-axis). Deep color indicates significantly dysregulated miRNAs.

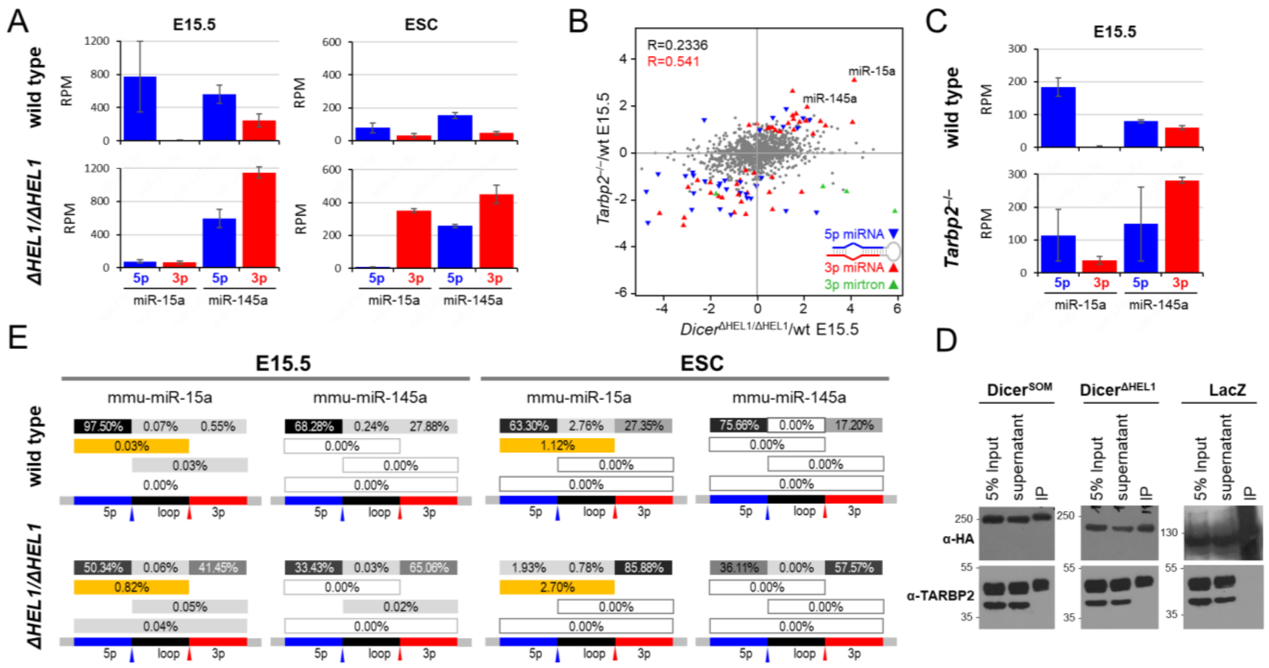


**Fig. S4. Strand selection bias and asymmetric cleavage.**

(**A**) Expression of miR-15a and miR-145a miRNAs in *Dicer*$^{\Delta HEL1/\Delta HEL1}$ mutants. Expression is shown in reads per million (RPM) of 19-25 nt RNA fragments. Error bars = SD. (**B**) Comparison of relative changes of miRNAs in *Tarbp2*$^{-/-}$ and *Dicer*$^{\Delta HEL1/\Delta HEL1}$ E15.5 embryos. Highlighted are miRNAs significantly differentially expressed in *Tarbp2*$^{-/-}$. Most miRNAs differentially expressed in *Tarbp2*$^{-/-}$ E15.5(*20*) showed changes in the same direction in *Dicer*$^{\Delta HEL1/\Delta HEL1}$ E15.5 embryos. (**C**) Expression of miR-15a and miR-145a miRNAs in *Tarbp2*$^{-/-}$ mutants. Expression is shown in reads per million (RPM) of 19-25 nt RNA fragments. Error bars = SD. (**D**) TARBP2 binds Dicer$^{\Delta HEL1}$. The western blots show TARBP2 presence in immunoprecipitates of Dicer$^{SOM}$ and Dicer$^{\Delta HEL1}$ isoforms. HA-tagged Dicer$^{SOM}$ or Dicer$^{\Delta HEL1}$ transiently expressed in NIH 3T3 cells were immunoprecipitated with α–HA antibody, and were analyzed by western blotting. The lower band in bottom western blots is a TARBP2 isoform, which does not interact with Dicer. (**E**) Products of asymmetric cleavage are detectable for pre-miR-15a but not for miR-145a in RNA-seq data from E15.5 and ESC samples. The schemes at the bottom of each panel represent genomic pre-miRNA sequence with 5p miRNA, loop and 3p miRNA from the left to the right. Dicer cleavage positions are depicted by blue (3′ end of 5p miRNA) and red (5′ end of 3p miRNA) arrowheads. Above are shown RNA fragments corresponding to pre-miRNA, mature miRNAs, the loop, and fragments cleaved only at the 3′ of 5p miRNA or 5′ of 3p miRNA. Numbers correspond to percentages observed in RNA-seq data from ESCs. Orange rectangles represent fragments of pre-miR-15a cleaved solely at the 5′ of 3p miRNA, dark grey rectangles depict most abundant fragments RNA-seq data mapping to the locus.

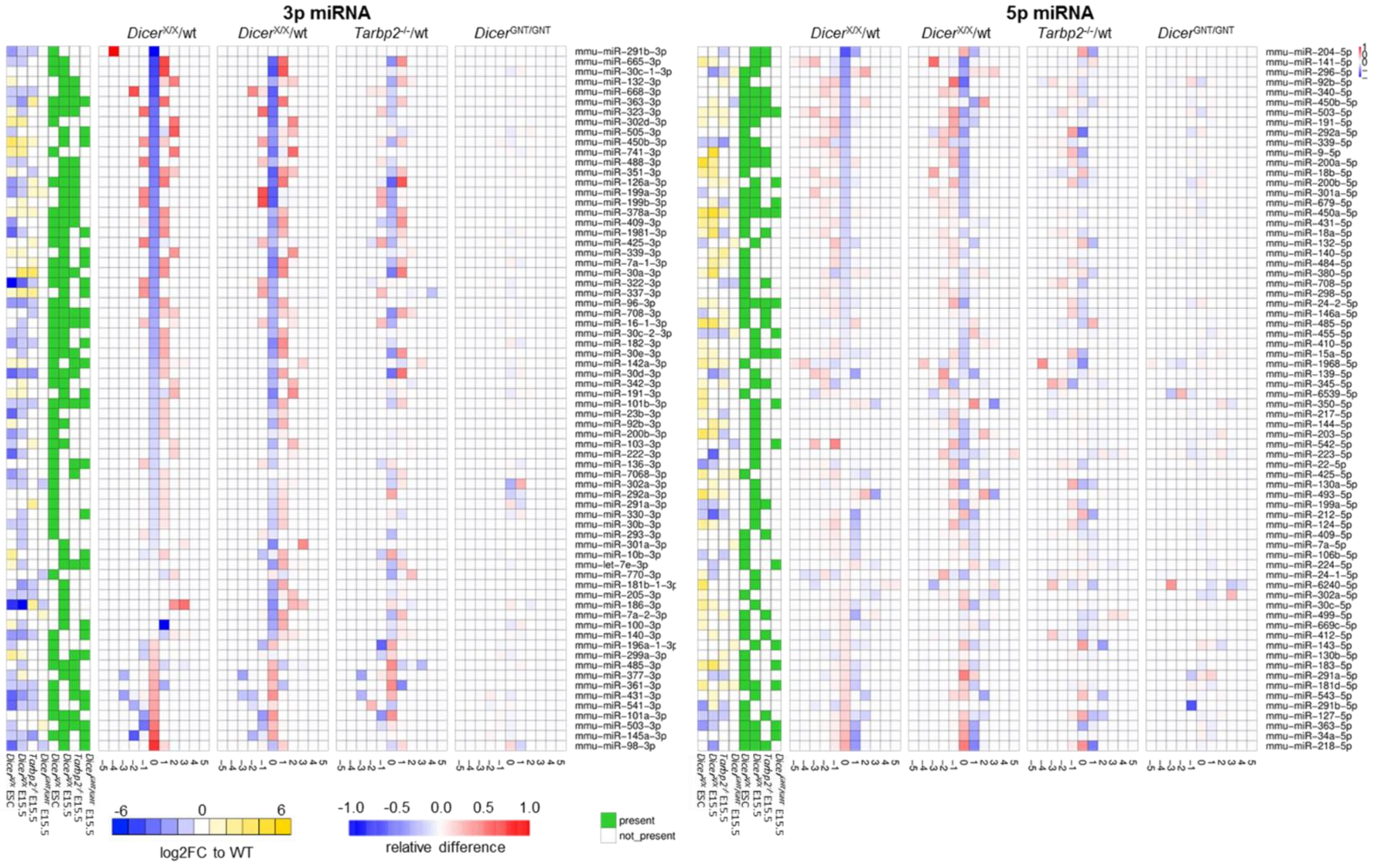

**Fig. S5. Cleavage fidelity in *Dicer*$^{\Delta HEL1/\Delta HEL1}$ mouse mutants.**

Heatmaps depict analysis of the 5′ 3p miRNA (left heatmap) and 3′ 5p miRNA (right heatmap) cleavage sites in 50 most affected miRNAs among all 3p miRNAs (>100 DESeq RPMs) in Dicer$^{\Delta HEL1/\Delta HEL1}$ E15.5 embryos and ESCs. At the center is the cleavage site. Each column of squares represents one nucleotide from the cleavage site in direction into the mature 3p miRNA (to the right) or upstream of it (to the left). Red-blue colors indicate relative changes in the 3p miRNA cleavage site when compared with the wild type sample.

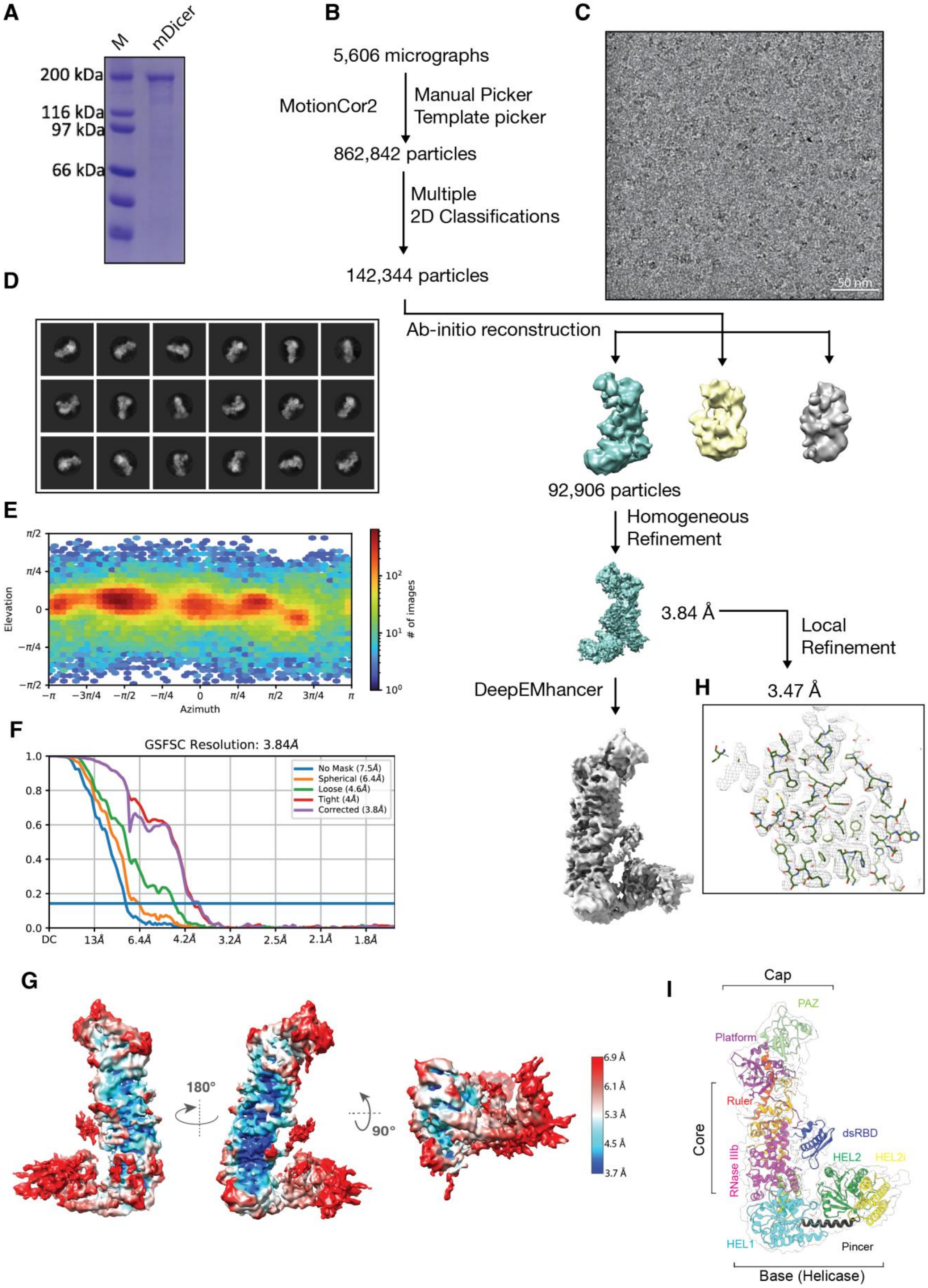
A
M
mDicer
200 kDa
116 kDa
97 kDa
66 kDa
B
5,606 micrographs
MotionCor2
Manual Picker
Template picker
862,842 particles
Multiple
2D Classifications
142,344 particles
C
50 nm
D
Ab-initio reconstruction
92,906 particles
Homogeneous
Refinement
3.84 Å
Local
Refinement
DeepEMhancer
H
3.47 Å
E
Elevation
Azimuth
### of images
F
GSFSC Resolution: 3.84Å
No Mask (7.5Å)
Spherical (6.4Å)
Loose (4.6Å)
Tight (4Å)
Corrected (3.8Å)
G
180°
90°
6.9 Å
6.1 Å
5.3 Å
4.5 Å
3.7 Å
I
Cap
PAZ
Platform
Ruler
Core
dsRBD
HEL2
HEL2i
RNase IIIb
HEL1
Pincer
Base (Helicase)

**Fig. S6. Purification and cryo-EM analyses of apo-Dicer.**

(**A**) SDS-page analysis of apo-Dicer. (**B**) Schematic outline of the image processing steps used to obtain the 3.84-Å-resolution cryo-EM reconstruction of apo-Dicer. 3D classes with no density of the helicase domain (due to inherent flexibility) were not used for the final reconstruction. (**C**) Representative cryo-EM micrograph of apo-Dicer. (**D**) Gallery of reference-free 2D class averages. (**E**) Heat map for distribution of particles for the final 3D reconstruction. (**F**) Gold standard FSC curves for apo-Dicer reconstructions. (**G**) Local resolution map of the final 3D reconstruction. (**H**) cryo-EM map from local refinement and fitting of coordinates. (**I**) Overall structure of mouse Dicer shown in ribbon representation.

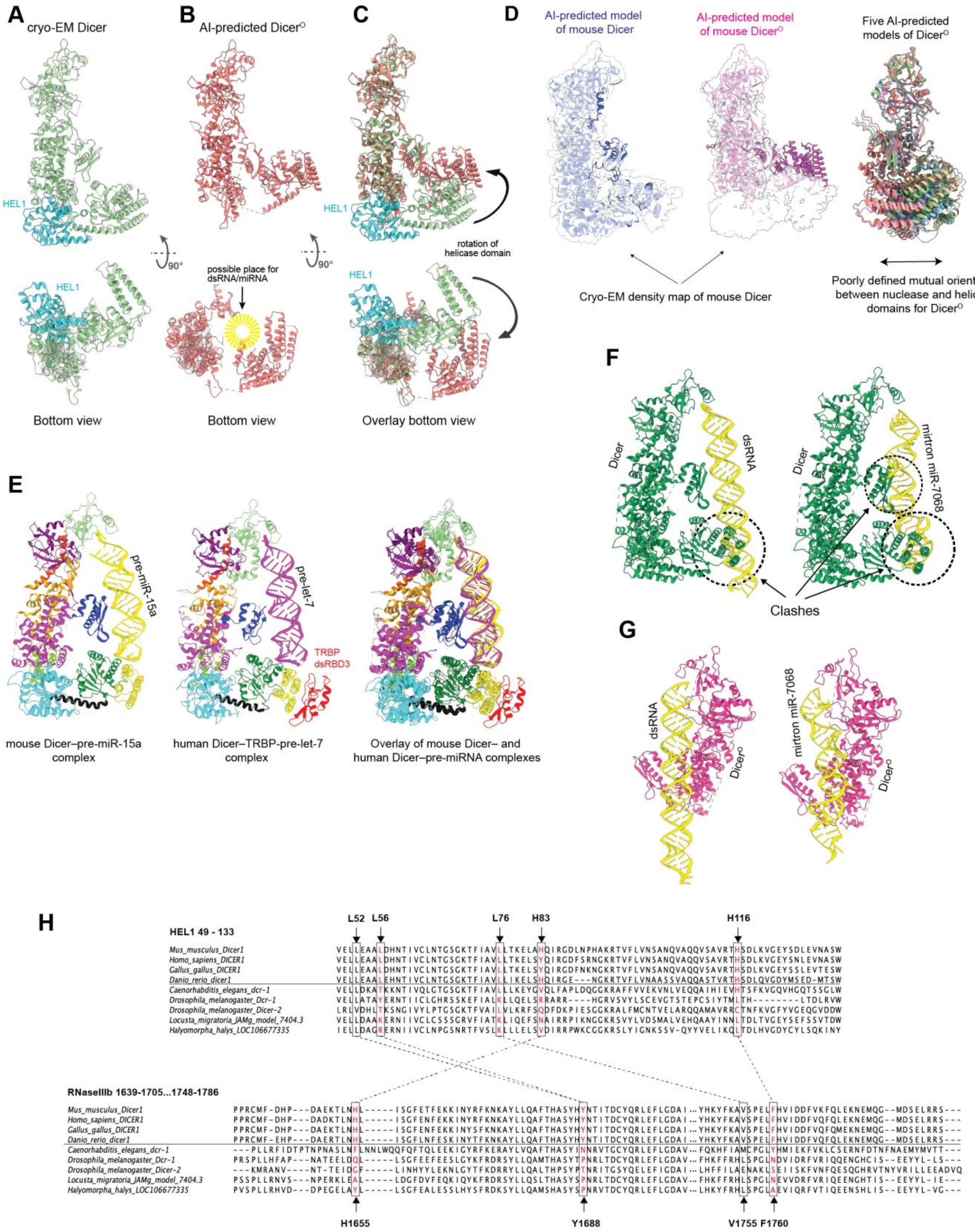
A
cryo-EM Dicer
HEL1
90°
Bottom view
B
AI-predicted Dicer$^{O}$
possible place for dsRNA/miRNA
Bottom view
C
HEL1
rotation of helicase domain
Overlay bottom view
D
AI-predicted model of mouse Dicer
AI-predicted model of mouse Dicer$^{O}$
Five AI-predicted models of Dicer$^{O}$
Cryo-EM density map of mouse Dicer
Poorly defined mutual orientation between nuclease and helicase domains for Dicer$^{O}$
E
pre-miR-15a
pre-let-7
TRBP dsRBD3
mouse Dicer–pre-miR-15a complex
human Dicer–TRBP-pre-let-7 complex
Overlay of mouse Dicer– and human Dicer–pre-miRNA complexes
F
Dicer
dsRNA
mirtron miR-7068
Clashes
G
dsRNA
Dicer$^{O}$
mirtron miR-7068
H
HEL1 49 - 133
L52 L56 L76 H83 H116
Mus_musculus_Dicer1
Homo_sapiens_DICER1
Gallus_gallus_DICER1
Danio_rerio_dicer1
Caenorhabditis_elegans_dcr-1
Drosophila_melanogaster_Dcr-1
Drosophila_melanogaster_Dicer-2
Locusta_migratoria_JAMg_model_7404.3
Halyomorpha_halys_LOC106677335
RNaseIIIb 1639-1705...1748-1786
H1655 Y1688 V1755 F1760

**Fig. S7. Comparison of Cryo-EM structures and AlphaFold models of Dicers.**

(**A**) The cryo-EM structure of mouse Dicer highlighting the position of HEL1 in the closed, inactive conformation of Dicer, shown in two orthogonal views. (**B**) AIrtifical intelligence (AI)-predicted model of Dicer$^{O}$, highlighting the open conformation of Dicer and possible accommodation of double-stranded RNA. (**C**) Superimposition of Dicer (green; HEL1 in magenta) and Dicer$^{O}$ (red), arrow indicates opening of helicase domain. (**D**) Validation of the AI-predicted Dicer model Dicer with the cryo-EM density map of Dicer (left). This supports reliability of artificial intelligence-based modeling, as AlphaFold correctly predicted the structure of full-length Dicer, including the correct mutual orientation between the core and helicase domain. The best AI-predicted model for Dicer$^{O}$ compared with the cryo-EM density map of Dicer (middle). Five Dicer$^{O}$ models (superimposed over the Dicer core and cap regions) by AlphaFold show that the models are predicted with a low confidence in the relative orientation between the helicase domain and the core, suggesting an increased flexibility in the absence of HEL1 (right). (**E**) The cryo-EM structure of murine Dicer (left) and human Dicer (middle) (PDB 5ZAL) in complex with pre-miRNAs in pre-dicing states shown in ribbon representation. Structural superimposition of murine Dicer–pre-miRNA and *Arabidopsis* DCL1–pre-miRNA complexes (right). (**F**) Compatibility of dsRNA/mirtrons binding with Dicer in the pre-dicing and dicing states. Dicer in the pre-dicing state cannot optimally bind long dsRNA or mirtrons due to steric hindrance (indicated by arrows). The model is build based of the cryo-EM structure of Dicer–pre-miR-15a complex, in which pre-miR-15a was replaced by a 42-bp dsRNA (left) or miR-7068 (right). (**G**) Dicer$^{O}$ in the dicing state can accommodate long dsRNA or mirtrons without steric hindrance. The model is constructed based of the cryo-EM structure of Dicer$^{O}$–pre-miR-15a, in which pre-miR-15a was replaced by a 42-bp dsRNA (left) or miR-7068 (right). (**H**) Multiple sequence alignments of HEL1 and RNase IIIb. Conserved residues in vertebrates depicted in red and their contacts in dotted lines.

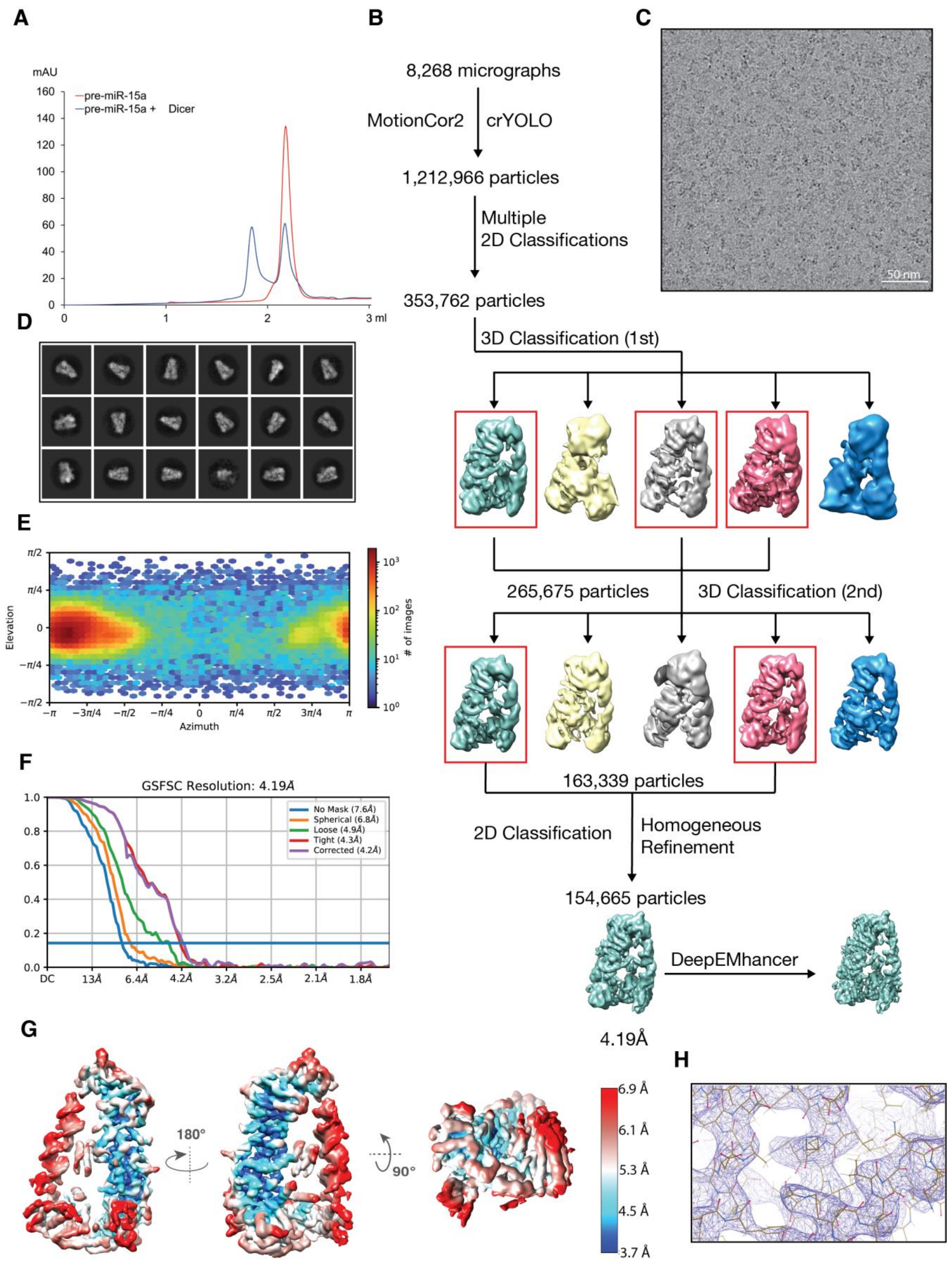
A
mAU
pre-miR-15a
pre-miR-15a + Dicer
3 ml
B
8,268 micrographs
MotionCor2
crYOLO
1,212,966 particles
Multiple
2D Classifications
353,762 particles
3D Classification (1st)
265,675 particles
3D Classification (2nd)
163,339 particles
2D Classification
Homogeneous
Refinement
154,665 particles
DeepEMhancer
4.19Å
C
50 nm
D
E
Elevation
Azimuth
### of images
F
GSFSC Resolution: 4.19Å
No Mask (7.6Å)
Spherical (6.8Å)
Loose (4.9Å)
Tight (4.3Å)
Corrected (4.2Å)
G
180°
90°
6.9 Å
6.1 Å
5.3 Å
4.5 Å
3.7 Å
H

**Fig. S8. Reconstitution and cryo-EM analyses of Dicer–pre-miR-15a complex.**

(**A**) Gel filtration analysis of Dicer–pre-miR-15a complex. (**B**) Schematic outline of the image processing steps used to obtain the 4.19-Å-resolution cryo-EM reconstruction of the Dicer–pre-miR-15a complex. (**C**) Representative cryo-EM micrograph of the Dicer–pre-miR-15a complex. (**D**) Gallery of reference-free 2D class averages. (**E**) Heat map for distribution of particles for the final 3D reconstruction. (**F**) Gold standard FSC curves for the Dicer–pre-miR-15a complex. (**G**) Local resolution map of the final 3D reconstruction. (**H**) cryo-EM map and fitting of coordinates.

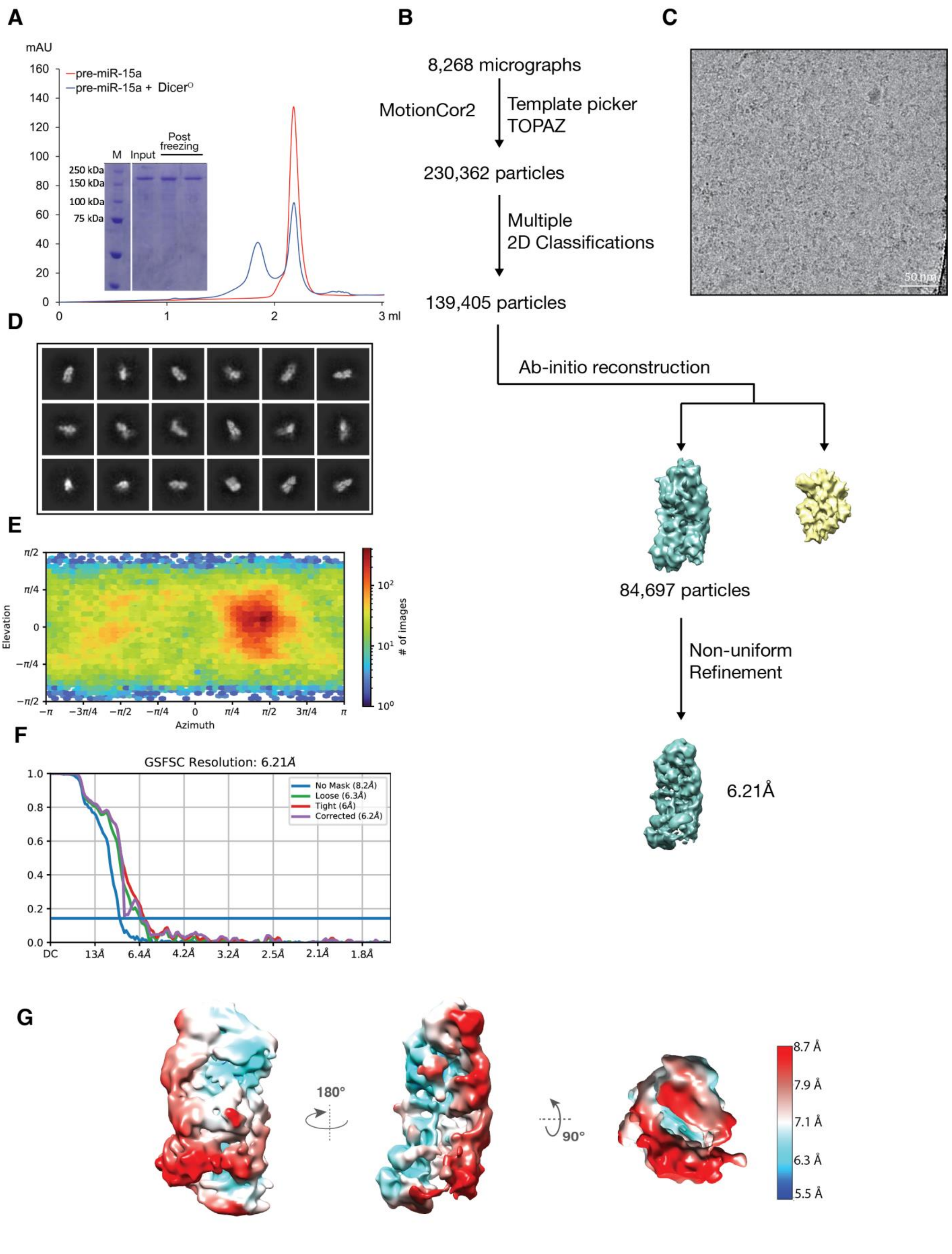
A
mAU
pre-miR-15a
pre-miR-15a + Dicer^O
M
Input
Post freezing
250 kDa
150 kDa
100 kDa
75 kDa
3 ml
B
8,268 micrographs
MotionCor2
Template picker
TOPAZ
230,362 particles
Multiple
2D Classifications
139,405 particles
Ab-initio reconstruction
84,697 particles
Non-uniform
Refinement
6.21Å
C
50 nm
D
E
Elevation
Azimuth
### of images
F
GSFSC Resolution: 6.21Å
No Mask (8.2Å)
Loose (6.3Å)
Tight (6Å)
Corrected (6.2Å)
G
180°
90°
8.7 Å
7.9 Å
7.1 Å
6.3 Å
5.5 Å

**Fig. S9. Reconstitution and cryo-EM analyses of Dicer$^{O}$–pre-miR-15a complex.**

(**A**) Gel filtration and SDS analyses of Dicer$^{O}$–pre-miR-15a complex. (**B**) Schematic outline of the image processing steps used to obtain the 6.21-Å-resolution cryo-EM reconstruction of the Dicer$^{O}$–pre-miR-15a complex. (**C**) Representative cryo-EM micrograph. (**D**) Gallery of reference-free 2D class averages. (**E**) Heat map for distribution of particles for the final 3D reconstruction. (**F**) Gold standard FSC curves for the Dicer$^{O}$–pre-miR-15a complex. (**G**) Local resolution map of the final 3D reconstruction.

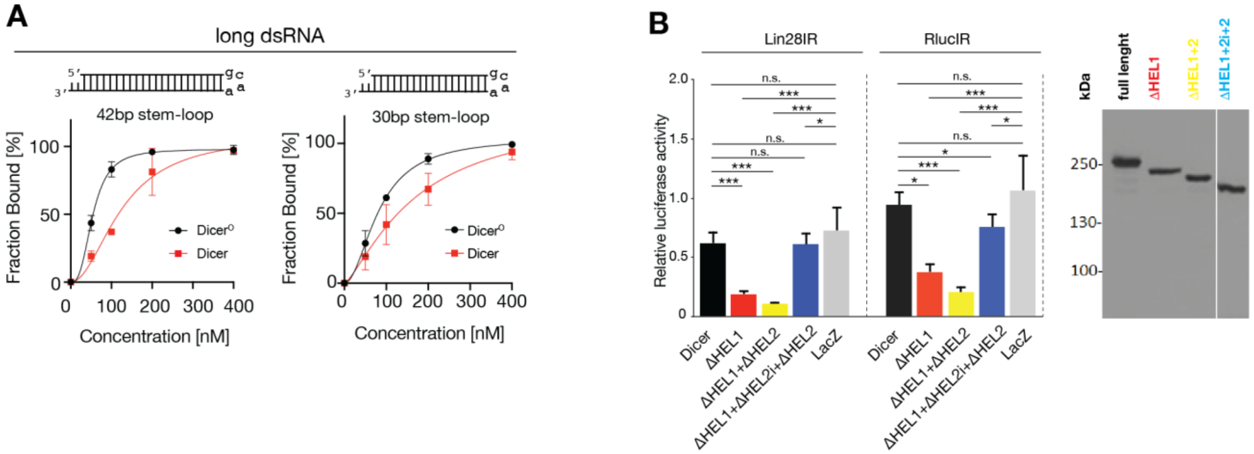


**Fig. S10. Characterization of Dicer and Dicer$^{O}$ *in vitro* and *in vivo*.**

(**A**) EMSA assays and their quantification for Dicer and Dicer$^{O}$ with dsRNAs capped by a gcaa-tetraloop, data points, mean ± SD (n=2-3). (**B**) RNA-mediated repression by different N-terminally truncated Dicer in *Pkr*$^{-/-}$ U-2 OS cells (left). Cells were co-transfected with plasmids expressing a human Dicer variant (or LacZ as a negative control), dsRNA (Lin28IR, RlucIR, or MosIR), a targeted *Renilla* luciferase reporter with complementary sequences to dsRNA from Lin28IR and RlucIR, and a non-targeted firefly luciferase. Expression of truncated Dicer proteins was confirmed by Western blotting (right). The y-scale shows the *Renilla* luciferase activity normalized to the non-targeted firefly luciferase activity, and non-specific effect of MosIR. Removing HEL1 or HEL1+HEL2 while retaining HEL2i yields similar robust RNAi effect with both dsRNAs. RNAi becomes weaker and more similar to the full-length Dicer when the deletion includes also HEL2i, the TARBP2 binding domain. Data points, mean ± SD (n=4-5; in triplicate transfections). *, $p < 0.05$; **, $p < 0.01$; ***, $p < 0.001$; $p > 0.05$, n.s. (non-significant).

#### Supplementary Tables

##### Table S1 miRNA expression in mutants

Table 1 is provided as a separate MS Excel file.

##### Table S2 Expression of host genes of most upregulated mirtrons in ESCs

| host gene | host gene id | mRNA baseMean | log2FC | pvalue | padj | mirtron |
|---|---|---|---|---|---|---|
| *Cherp* | ENSMUSG00000052488.7 | 617.8 | 0.845 | 0.004 | 0.377 | mmu-miR-7068 |
| *Dbn1* | ENSMUSG00000034675.17 | 52.0 | -0.304 | 0.542 | 1.000 | mmu-miR-6944 |
| *Arap3* | ENSMUSG00000024451.8 | 10.4 | 0.614 | 0.358 | 1.000 | mmu-miR-6981 |
| *Fbrs* | ENSMUSG00000042423.9 | 197.3 | 0.219 | 0.520 | 1.000 | mmu-miR-7060 |
| *Dennd6b* | ENSMUSG00000015377.9 | 163.9 | 0.105 | 0.791 | 1.000 | mmu-miR-6958 |
| *Gfra4* | ENSMUSG00000027316.15 | 11.4 | -0.720 | 0.280 | 1.000 | mmu-miR-6973b |
| *Fbxw9* | ENSMUSG00000008167.14 | 711.2 | 0.223 | 0.536 | 1.000 | mmu-miR-7070 |
| *Hspg2* | ENSMUSG00000028763.17 | 2418.0 | -0.348 | 0.264 | 1.000 | mmu-miR-7018 |
| *Nav1* | ENSMUSG00000009418.15 | 451.2 | 0.807 | 0.006 | 0.499 | mmu-miR-1231 |
| *Atp2b4* | ENSMUSG00000026463.17 | 275.5 | -1.178 | 0.003 | 0.351 | mmu-miR-6903 |
| *Arhgef17* | ENSMUSG00000032875.8 | 301.1 | -0.549 | 0.085 | 1.000 | mmu-miR-3102 |
| *Etfb* | ENSMUSG00000004610.4 | 666.6 | -0.477 | 0.121 | 1.000 | mmu-miR-7051 |
| *Ptprs* | ENSMUSG00000013236.17 | 602.1 | 0.012 | 0.971 | 1.000 | mmu-miR-6977 |
| *Rrp1* | ENSMUSG00000061032.9 | 1706.2 | -0.101 | 0.757 | 1.000 | mmu-miR-6907 |
| *Myh3* | ENSMUSG00000020908.14 | 27.6 | -0.389 | 0.473 | 1.000 | mmu-miR-6923 |
| *Baiap3* | ENSMUSG00000047507.12 | 4.4 | -0.270 | 0.733 | 1.000 | mmu-miR-3547 |
| *Ciao3* | ENSMUSG00000002280.10 | 367.9 | -0.313 | 0.296 | 1.000 | mmu-miR-6966 |
| *Hip1r* | ENSMUSG00000000915.15 | 189.2 | 0.241 | 0.499 | 1.000 | mmu-miR-7032 |
| *Farsa* | ENSMUSG00000003808.18 | 1774.8 | 0.373 | 0.228 | 1.000 | mmu-miR-7069 |
| *Mst1* | ENSMUSG00000032591.15 | 93.0 | -0.334 | 0.458 | 1.000 | mmu-miR-7088 |
| *Ap2a2* | ENSMUSG00000002957.11 | 2406.3 | 0.011 | 0.972 | 1.000 | mmu-miR-7063 |
| *Dnase1l1* | ENSMUSG00000019088.13 | 27.5 | -0.661 | 0.207 | 1.000 | mmu-miR-7091 |

#### Table S3 Asymmetric cleavage of miRNAs

Shown are frequencies of specific miRNA fragments in RNA-sequencing data from *Dicer*$^{\Delta HEL1/\Delta HEL1}$ ESCs. The fragment miR-5p+loop is produced by asymmetric cleave at the 5' end of 3p miRNA.

| miRNA | miR-5p | loop | mR-3p | **miR-5p+loop** | miR-3p+loop |
|---|---|---|---|---|---|
| mmu-miR-7041 | 0.0000 | 0.0000 | 0.6667 | **0.3333** | 0.0000 |
| mmu-miR-7067 | 0.0000 | 0.0000 | 0.3333 | **0.3333** | 0.0000 |
| mmu-miR-667 | 0.5358 | 0.0037 | 0.0838 | **0.3322** | 0.0000 |
| mmu-miR-31 | 0.5807 | 0.0000 | 0.0519 | **0.2948** | 0.0031 |
| mmu-miR-29c | 0.1667 | 0.0000 | 0.1667 | **0.2222** | 0.0000 |
| mmu-miR-101b | 0.0000 | 0.0000 | 0.8350 | **0.1456** | 0.0012 |
| mmu-miR-465b | 0.5317 | 0.0000 | 0.3394 | **0.1106** | 0.0000 |
| mmu-miR-465b | 0.5317 | 0.0000 | 0.3394 | **0.1106** | 0.0000 |
| mmu-miR-539 | 0.2650 | 0.2222 | 0.0513 | **0.0769** | 0.0256 |
| mmu-miR-154 | 0.2100 | 0.0284 | 0.6601 | **0.0664** | 0.0002 |
| mmu-miR-3070 | 0.2405 | 0.0000 | 0.6881 | **0.0476** | 0.0000 |
| mmu-miR-465c | 0.5198 | 0.0000 | 0.4097 | **0.0407** | 0.0000 |
| mmu-miR-465c | 0.5198 | 0.0000 | 0.4097 | **0.0407** | 0.0000 |
| mmu-miR-142a | 0.3436 | 0.0082 | 0.3154 | **0.0402** | 0.0000 |
| mmu-miR-377 | 0.5106 | 0.0000 | 0.4012 | **0.0359** | 0.0000 |
| mmu-miR-367 | 0.0000 | 0.0827 | 0.7694 | **0.0351** | 0.0000 |
| mmu-miR-878 | 0.6171 | 0.0000 | 0.2904 | **0.0334** | 0.0000 |
| mmu-miR-293 | 0.2776 | 0.0000 | 0.6612 | **0.0291** | 0.0001 |
| **mmu-miR-15a** | **0.0193** | **0.0078** | **0.8588** | **0.0270** | **0.0000** |
| mmu-miR-141 | 0.1689 | 0.1947 | 0.3683 | **0.0264** | 0.0034 |
| mmu-miR-324 | 0.0620 | 0.0361 | 0.8431 | **0.0250** | 0.0000 |
| mmu-miR-883b | 0.5185 | 0.0000 | 0.4577 | **0.0238** | 0.0000 |
| mmu-miR-376b | 0.0291 | 0.0173 | 0.9008 | **0.0235** | 0.0000 |
| mmu-miR-485 | 0.3695 | 0.0001 | 0.5528 | **0.0214** | 0.0146 |
| mmu-miR-20b | 0.9475 | 0.0006 | 0.0089 | **0.0199** | 0.0004 |
| mmu-miR-743b | 0.0394 | 0.0000 | 0.9235 | **0.0185** | 0.0000 |
| mmu-miR-665 | 0.0400 | 0.0000 | 0.8164 | **0.0176** | 0.0820 |
| mmu-miR-679 | 0.7761 | 0.0000 | 0.0568 | **0.0167** | 0.0000 |
| mmu-miR-465a | 0.0921 | 0.0000 | 0.8556 | **0.0143** | 0.0057 |
| mmu-miR-181c | 0.0812 | 0.0000 | 0.8520 | **0.0140** | 0.0000 |
| mmu-miR-188 | 0.9352 | 0.0139 | 0.0000 | **0.0139** | 0.0000 |
| mmu-miR-411 | 0.7924 | 0.0023 | 0.1631 | **0.0139** | 0.0000 |
| mmu-miR-93 | 0.9053 | 0.0019 | 0.0396 | **0.0135** | 0.0000 |
| mmu-miR-301a | 0.5232 | 0.0067 | 0.1228 | **0.0133** | 0.0000 |
| mmu-miR-362 | 0.4786 | 0.0000 | 0.4550 | **0.0130** | 0.0000 |
| mmu-miR-341 | 0.0241 | 0.0000 | 0.8388 | **0.0121** | 0.0003 |
| mmu-miR-380 | 0.3384 | 0.0000 | 0.5523 | **0.0115** | 0.0000 |
| mmu-miR-211 | 0.8401 | 0.0000 | 0.1078 | **0.0109** | 0.0021 |
| mmu-miR-29a | 0.0085 | 0.0000 | 0.9267 | **0.0096** | 0.0000 |
| mmu-miR-677 | 0.2121 | 0.0000 | 0.0004 | **0.0089** | 0.0025 |

#### Table S4 Cryo-EM data collection and refinement statistics

| | | | |
|---|---|---|---|
| **Instrument** | | | |
| Microscope | FEI Titan Krios | | |
| Camera | Gatan K2 Summit direct electron camera (counting mode) | | |
| **Data collection** | | | |
| **Sample** | **Dicer** | **Dicer–pre-miR-15a** | **Dicer$^{O}$–pre-miR-15a** |
| Voltage (kV) | 300 | 300 | 300 |
| Electron dose ($e^-$/Å$^2$) | 55.0 | 55.0 | 55.0 |
| Defocus range (μm) | -0.8 to -3.5 | -1 to -3.5 | -1 to -3.5 |
| Pixel size (Å) | 0.828 | 0.828 | 0.828 |
| Movies collected | 6,354 | 16,601 | 9,956 |
| Initial particles | 862,842 | 1,212,966 | 230,362 |
| Final particles | 92,906 | 154,665 | 84,697 |
| **Model composition** | | | |
| Protein residues | 1234 | 1281 | 705 |
| **Refinement** | | | |
| Map resolution (Å) | 3.84 | 4.19 | 6.21 |
| FSC threshold | 0.143 | 0.143 | 0.143 |
| Combined map resolution range (Å) | 3.47 – 4.18 | - | - |
| CC mask | 0.609 | 0.731 | 0.524 |
| CC volume | 0.617 | 0.739 | 0.501 |
| CC peaks | 0.536 | 0.649 | 0.390 |
| Map sharpening *B*-factor (Å$^2$) | 87.8 | 134.8 | 451.4 |
| **R.m.s deviations** | | | |
| Bond lengths (Å) | 0.004 | 0.004 | 0.013 |
| Bond angles (°) | 0.780 | 0.798 | 1.917 |
| **Validation** | | | |
| MolProbity score | 1.12 | 1.07 | 0.78 |
| Clashscore | 1.46 | 1.57 | 0.92 |
| Poor rotamers (%) | 0.92 | 0.18 | 0.64 |
| **Ramachandran plot** | | | |
| Favored (%) | 96.44 | 97.07 | 98.70 |
| Allowed (%) | 3.56 | 2.93 | 1.30 |
| Disallowed (%) | 0.00 | 0.00 | 0.00 |

#### Table S5 RNA-seq libraries

| stage | type | genotype | library name | note |
|---|---|---|---|---|
| ESC | small RNA | *Dicer$^{wt/wt}$* | s_ESC_WT+MosIR_RS7.1 | transfected with MosIR |
| ESC | small RNA | *Dicer$^{wt/wt}$* | s_ESC_WT+MosIR_RS7.2 | transfected with MosIR |
| ESC | small RNA | *Dicer$^{wt/wt}$* | s_ESC_WT+MosIR_RS7.3 | transfected with MosIR |
| ESC | small RNA | *Dicer$^{\Delta HEL1/\Delta HEL1}$* | s_ESC_XHOM+MosIR_RS10.1 | transfected with MosIR |
| ESC | small RNA | *Dicer$^{\Delta HEL1/\Delta HEL1}$* | s_ESC_XHOM+MosIR_RS10.2 | transfected with MosIR |
| ESC | small RNA | *Dicer$^{\Delta HEL1/\Delta HEL1}$* | s_ESC_XHOM+MosIR_RS10.3 | transfected with MosIR |
| E15.5 | small RNA | *Dicer$^{wt/wt}$* | s_E15.5_WT_1 | |
| E15.5 | small RNA | *Dicer$^{wt/wt}$* | s_E15.5_WT_6 | |
| E15.5 | small RNA | *Dicer$^{wt/wt}$* | s_E15.5_WT_8B | |
| E15.5 | small RNA | *Dicer$^{\Delta HEL1/\Delta HEL1}$* | s_E15.5_XHOM_10B_r2 | |
| E15.5 | small RNA | *Dicer$^{\Delta HEL1/\Delta HEL1}$* | s_E15.5_XHOM_2 | |
| E15.5 | small RNA | *Dicer$^{\Delta HEL1/\Delta HEL1}$* | s_E15.5_XHOM_3B | |
| E15.5 | small RNA | *Dicer$^{\Delta HEL1/\Delta HEL1}$* | s_E15.5_XHOM_4 | |
| E15.5 | small RNA | *Dicer$^{\Delta HEL1/\Delta HEL1}$* | s_E15.5_XHOM_7B | |
| E15.5 | small RNA | *Dicer$^{wt/wt}$* | s_E15.5_WT_11 | |
| E15.5 | small RNA | *Dicer$^{wt/wt}$* | s_E15.5_WT_14 | |
| E15.5 | small RNA | *Dicer$^{wt/wt}$* | s_E15.5_WT_16 | |
| E15.5 | small RNA | *Dicer$^{GNT/GNT}$* | s_E15.5_GNTHOM_3 | |
| E15.5 | small RNA | *Dicer$^{GNT/GNT}$* | s_E15.5_GNTHOM_4 | |
| E15.5 | small RNA | *Dicer$^{GNT/GNT}$* | s_E15.5_GNTHOM_9 | |
| E15.5 | small RNA | *Tarbp2$^{+/+}$* | SRS2781156 B6T2-65Tarbp2_WT_1 | PRJNA423238 SRP127346 |
| E15.5 | small RNA | *Tarbp2$^{+/+}$* | SRS2781151 B6T2-65Tarbp2_WT_2 | PRJNA423238 SRP127346 |
| E15.5 | small RNA | *Tarbp2$^{+/+}$* | SRS2781150B6T2-65Tarbp2_WT_3 | PRJNA423238 SRP127346 |
| E15.5 | small RNA | *Tarbp2$^{-/-}$* | SRS2781155 B6T2-54Tarbp2_Mut_1 | PRJNA423238 SRP127346 |
| E15.5 | small RNA | *Tarbp2$^{-/-}$* | SRS2781154 B6T2-60Tarbp2_Mut_2 | PRJNA423238 SRP127346 |
| E15.5 | small RNA | *Tarbp2$^{-/-}$* | SRS2781157 B6T2-90Tarbp2_Mut_3 | PRJNA423238 SRP127346 |

#### Table S6. Primers for mouse Dicer constructs and synthetic RNAs.

| **Oligonucleotides** | | |
|---|---|---|
| 2xFLAG_F:<br>cacgacatcgactacaaggacgacgacgacaagTGAAGCGGCCGCTTCCCT | Sigma-Aldrich | - |
| 2xFLAG_Rev:<br>gtccttgtagtcaccgtcgtggtccttgtagtcGCTATTGGGAACCTGAGGTTGATTAGC | Sigma-Aldrich | - |
| dHEL1_F:<br>GGGCTTTATGAAAGACTGC | Sigma-Aldrich | - |
| dHEL1_R:<br>TTGCAAAGCAGGGCTTTT | Sigma-Aldrich | - |
| dDExD_F:<br>AACACGGCCATTGGACAC | Sigma-Aldrich | - |
| dDExD_R:<br>see dHEL1_R | Sigma-Aldrich | - |
| dHEL2_F:<br>see dDExD_F | Sigma-Aldrich | - |
| dHEL2_R:<br>TAAGACAACTGCTGTGTATCTTC | Sigma-Aldrich | - |
| Deletion confirmation - forward primer:<br>CCGTTCATTTCCCAGCCTGT | Sigma-Aldrich | - |
| Deletion confirmation - reverse primer:<br>AAAACAGCCCAATTCCTTGCC | Sigma-Aldrich | - |
| Twin-HA-TEV_Fwd:<br>ATCTACGGATCCACCATGGTATGGAGCCATCCTCAATTTGAAAAGGGTGGCG<br>GGTCCGGCGGTGGGTCTGGCGGTAGCGCTTGGTCCCACCCCCAGTTCG | Sigma-Aldrich | - |
| Twin-HA-TEV_Rev:<br>GTAGATGTCGACCAGGCCCTGAAAATACAGGTTTTCGGTACCAGCGTAATCT<br>GGAACATCGTATGGGTAGTCACCCTTCTCGAACTGGGGGTGGGACCAA | Sigma-Aldrich | - |
| C_TEV-FLAG-His_Fwd:<br>TCTACAGCGGCCGCGGCGAGAATCTCTACTTCCAAGGCGCTAGCGACTATAA<br>GGACCACGACGGAGACTA | Sigma-Aldrich | - |
| C_TEV-FLAG-His_Rev:<br>GTAGATAAGCTTAGTGATGGTGATGGTGATGGTGGTGGGACCCATCATGATC<br>CTTGTAGTCTCCGTCGTGGTCCTT | Sigma-Aldrich | - |
| mDicer_SalI_Fwd:<br>ATGTCGACGCAGGCCTGCAGCTCATGACCCC | Sigma-Aldrich | - |
| mDicerO_SalI_Fwd:<br>CAGTCGACAGCCGTGATACAGAAGTATACAC | Sigma-Aldrich | - |
| mDicer-NotI_Rev:<br>ATGCGGCCGCTGTTAGGAACCTGAGGCTGGTTAGC | Sigma-Aldrich | - |
| mDicer E1560A Forward:<br>GCTGACAAGAGCATAGCGGACTGTGTTGCTGCACTGCTGGGCTGCTACTTAA<br>CCAGC | Sigma-Aldrich | - |
| mDicer E1560A Reverse:<br>GCTGGTTAAGTAGCAGCCCAGCAGTGCAGCAACACAGTCCGCTATGCTCTTG<br>TCAGC | Sigma-Aldrich | - |
| mDicer E1807A Forward:<br>CAAGGCCATGGGGGACATTTTTGCATCTCTTGCTGGTGCCATTTATAT | Sigma-Aldrich | - |
| mDicer E1807A Reverse:<br>ATATAAATGGCACCAGCAAGAGATGCAAAAATGTCCCCCATGGCCTTG | Sigma-Aldrich | - |
| sgRNA targeting intron 2, No.1:<br>GAGATGAGTCCTATAAAGGGG | Sigma-Aldrich | - |

| | | |
|---|---|---|
| sgRNA targeting intron 2, No.2:<br>CCCCTCTGTCTCCTAAACTGC | Sigma-Aldrich | - |
| sgRNA targeting intron 2, No.3:<br>ACGGGAAGAAGAAATGGCTGG | Sigma-Aldrich | - |
| sgRNA targeting intron 8, No.1:<br>GCCATCTAGATATACAGGAGG | Sigma-Aldrich | - |
| sgRNA targeting intron 8, No.2:<br>CCTTACCCTTCCACACGTCAC | Sigma-Aldrich | - |
| pre-miR-15a:<br>UAGCAGCACAUAAUGGUUUGUGGAUGUUGAAAAGGUGCAGGCCAUACUGUGCUGCCUCA | Sigma-Aldrich | - |
| pre-miR-145a:<br>GUCCAGUUUUCCCAGGAAUCCCUUGGAUGCUAAGAUGGGGAUUCCUGGAAAUACUGUUCUUG | Sigma-Aldrich | - |
| pre-miR-7068:<br>GUGAGGCUCAGUAUGGGGUGGGGGUGUCGUCGCCUGCCCGACUGACCACCCACUCACCCUGGACUGACUCUCAG | Sigma-Aldrich | - |
| 30bp stem-loop:<br>AGAGGAGAGGGACAAUCAUAAAGGCCACUCGCAAGAGUGGCCUUUAUGAUUGUCCCUCUCCUCUUU | Sigma-Aldrich | - |
| 42bp stem-loop:<br>AGAGGAGAGGGACAAUAGAGGAGAGGGACAAUCAUAAAGGCCGCAAGGCCUUUAUGAUUGUCCCUCUCCUCUAUUGUCCCUCUCCUCUUU | Sigma-Aldrich | - |
| mDcr_i1_Fwd:<br>GATATAACCAGCTCAAGTGTTTGC | Sigma-Aldrich | genotyping (1st round of nested PCR) |
| mDcr_i7_Rev:<br>GAGCAAAAAGTTCATCAGGAACC | Sigma-Aldrich | genotyping (1st round of nested PCR) |
| mDcr_i1_Fwd2:<br>GCCTGGTTGGGTATAGACTGCTTG | Sigma-Aldrich | genotyping (2nd round of nested PCR) |
| mDcr_i7_Rev2:<br>CAGAGGGCTAGAGCATACAAACAC | Sigma-Aldrich | genotyping (2nd round of nested PCR) |
| Dicer_26720F:<br>CAAGCCCGCCTCTTCTGATT | Sigma-Aldrich | DicerGNT genotyping |
| Dicer_28976R:<br>ATGGCACGAATGACTGAACC | Sigma-Aldrich | DicerGNT genotyping |
| DQCH_30860F<br>CAGGTCTCATCTGCCAAGGT | Sigma-Aldrich | DicerDQCH genotyping |
| DQCH_33000R:<br>TGGAAGCAAGGCTTAGGAAA | Sigma-Aldrich | DicerDQCH genotyping |
